# Poly(ADP-ribosyl)ation associated changes in CTCF-chromatin binding and gene expression in breast cells

**DOI:** 10.1101/175448

**Authors:** Ioanna Pavlaki, France Docquier, Igor Chernukhin, Georgia Kita, Svetlana Gretton, Christopher T. Clarkson, Vladimir B. Teif, Elena Klenova

**Affiliations:** University of Essex, School of Biological Sciences, Wivenhoe Park, Colchester, Essex CO4 3SQ, UK; Current address: Department of Biology and Biochemistry, University of Bath, Claverton Down, Bath, BA2 7AY, UK; Current address: Department of Medical Genetics, Academic Laboratory of Medical Genetics, University of Cambridge, Box 238, Lv 6 Addenbrooke’s Treatment Centre, Cambridge Biomedical Campus, Cambridge CB2 0QQ, UK; Current address: Cancer Research UK Cambridge Institute, University of Cambridge, Li Ka Shing Centre, Robinson Way, Cambridge CB2 0RE, UK; Current address: Novartis (Hellas) S.A.C.I., Oncology Medical Affairs, National Road No 1, 12^th^, 144 51-Metamorphosis, Athens, Greece; Current address: University of Suffolk, Waterfront Building, Neptune Quay, Ipswich, IP4 1QJ, UK

## Abstract

CTCF is an evolutionarily conserved and ubiquitously expressed architectural protein regulating a plethora of cellular functions via different molecular mechanisms. CTCF can undergo a number of post-translational modifications which change its properties and functions. One such modifications linked to cancer is poly(ADP-ribosyl)ation (PARylation). The highly PARylated CTCF form has an apparent molecular mass of 180 kDa (referred to as CTCF180), which can be distinguished from hypo- and non-PARylated CTCF with the apparent molecular mass of 130 kDa (referred to as CTCF130). The existing data accumulated so far have been mainly related to CTCF130. However, the properties of CTCF180 are not well understood despite its abundance in a number of primary tissues. In this study we performed ChIP-seq and RNA-seq analyses in human breast cells 226LDM, which display predominantly CTCF130 when proliferating, but CTCF180 upon cell cycle arrest. We observed that in the arrested cells the majority of sites lost CTCF, whereas fewer sites gained CTCF or remain bound (i.e. common sites). The classical CTCF binding motif was found in the lost and common, but not in the gained sites. The changes in CTCF occupancies in the lost and common sites were associated with increased chromatin densities and altered expression from the neighboring genes. Based on these results we propose a model integrating the CTCF130/180 transition with CTCF-DNA binding and gene expression changes. This study also issues an important cautionary note concerning the design and interpretation of any experiments using cells and tissues where CTCF180 may be present.

## 1. INTRODUCTION

The CCCTC-binding factor (CTCF) is an evolutionarily conserved and ubiquitous chromatin protein that regulates 3D genome architecture and participates in multiple cellular functions including transcriptional activation, silencing, insulation, mediation of long range chromatin interactions and others [1–8]. Significant efforts are currently devoted to the investigation of molecular mechanisms of CTCF functioning in normal cells and disease using new generations of high-throughput sequencing [9–11]. This question is particularly important because CTCF binds to numerous sites of unclear function in the human genome, and some of these binding sites differ between different cells of the same organism [6, 9, 10, 12, 13].

Post-translational modifications of chromatin proteins (histones, transcription factors and others) are known to play an important role in differential protein binding in chromatin. Poly(ADP-ribosyl)ation (PARylation) is one of such modifications performed by poly(ADP-ribose) polymerases (PARPs) [14, 15]. Phylogenetically ancient PARylation is involved in the regulation of numerous cellular functions, such as DNA repair, replication, transcription, translation, telomere maintenance and chromatin remodeling [16–19]. A growing body of evidence demonstrates the link between CTCF PARylation and its biological functions. For example, the insulator and transcription factor functions of CTCF have been found to be regulated by PARylation [20, 21]. The effect of CTCF PARylation is important in DNA damage response [22]. A number of studies reported direct interaction between CTCF and poly(ADP-ribose) polymerase 1 (PARP1), as well as their co-localization in chromatin [23–25]. Furthermore, PARP1 and CTCF have been found to regulate the transition between active and repressed chromatin at the lamina [26]. A highly PARylated form of CTCF is represented by a protein with an apparent molecular mass 180 kDa (CTCF180), whereas the commonly observed CTCF130, is hypo- or non-PARylated. CTCF130 has been found in many immortalized cell lines and cancer tissues [23, 27–29]. Interestingly, only CTCF180 was detected in normal breast tissues, whereas both CTCF130 and CTCF180 were present in breast tumours [29]. Usually CTCF130 is associated with cell proliferation, whereas CTCF180 is characteristic for non-proliferating cells of different types. The latter include cells from healthy breast tissues with very low proliferative index [29], cells with induced cell cycle arrest, DNA damage [29], senescence [30] or apoptosis [28, 29]. Currently all existing information regarding the binding characteristics of CTCF has been mined from the experimental data obtained for CTCF130, but not CTCF180. It is not known whether the sets of targets for CTCF130 and CTCF180 are the same, completely different or overlap, and how binding of different forms of CTCF may be associated with alteration in gene expression. One of the reasons for this is that it is difficult to distinguish between CTCF130 and CTCF180 is the absence of an antibody specifically recognising CTCF180. All existing anti-CTCF antibodies detect either only CTCF130 or both CTCF130 and CTCF180. Furthermore, the antibody property differs from batch to batch even for the same commercial vendor, and in order to select the antibody with well-defined properties one has to perform screening of several batches, e.g. using Western blot assays.

In the present study we distinguished between CTCF130 and CTCF180 binding using a specific biological system: the immortalized human luminal breast cell line, 226LDM, which contains mainly non-PARylated CTCF (CTCF130) in the proliferating cell state, and mainly highly PARylated CTCF (CTCF180) upon cell cycle arrest with hydroxyurea (HU) and nocodazole (NO) [29]. We have previously proved that the form of CTCF migrating in the gel with the apparent molecular mass 180 kDa was PARylated (CTCF180) because (i) it could be generated from the recombinant CTCF by *in vitro* PARylation, (ii) it immunoprecipitated using anti-PAR antibodies and (iii) it contained peptides specific for CTCF [27]. Additional data presented below also confirm that CTCF180 can be detected exclusively in normal primary breast tissues, whereas both CTCF180 and CTCF130 are present in MCF7 (cancer) and 226LDM (immortalized) breast cells. Thus, the 226LDM cell model provides us with the unique opportunity to study both CTCF forms even in the absence of a specific antibody against CTCF180. Using this technique we aimed here to analyse the genomic targets for CTCF130 and CTCF180 in two functional states of 226LDM cells and connect these to the changes of chromatin states and gene expression.

## 2. MATERIAL AND METHODS

### 2.1. Cell Culture

226LDM cells, derived from human luminal breast cells, were propagated and cell cycle arrested as previously described [29]. In brief, cells were seeded in flasks and grown in DMEM/F-12 (PAA) supplemented with 5 μg / ml insulin, 1 μg / ml hydrocortisone, 20 ng / ml epidermal growth factor, 20 ng / ml cholera toxin (all from Sigma), 10 % fetal bovine serum (FBS) (Biosera), and 50 μg / ml gentamicin (Life Technologies-Invitrogen) at 37°C and 5 % CO_2_. To achieve full transition from CTCF130 to CTCF180 upon cell cycle arrest, the treatment conditions were further optimised. In particular, 226LDM cells were exposed to 100 mM hydroxyurea for 24 h followed by 1 h of complete medium, and a further 24 h with 500 ng / ml nocodazole (SIGMA). Cells in suspension were then harvested and assessed by Western blot assay to confirm complete transition (i.e. the presence of CTCF180 only and disappearance of CTCF130). These cells were 79% viable according to Countess^®^ automated cell counter (Life Sciences, USA) and arrested in the S- and G2/M-phases [29]. Untreated adherent proliferating 226LDM cells were used as control.

### 2.2. Immunoblotting

The endogenous protein levels of CTCF were observed by SDS-PAGE/ western blot analysis [31, 32] in whole cell lysates of 226LDM cells from the control and treated populations using a polyclonal anti-CTCF antibody (Millipore, 07-729, lot # JBC1903613, pre-screened with the lysates from breast normal and tumour tissues to detect both, CTCF130 and CTCF180). Anti-tubulin specific antibody (SIGMA, T5168) was used as a loading control. Chemiluminescence detection was performed with the Fusion FX7 gel documentation system (PeqLab) and the UptiLight (Interchim) reagents according to the manufacturer’s instructions.

### 2.3. Protein immunoprecipitation (IP)

CTCF IP was performed in 226LDM cells, using anti-CTCF antibody [33]. 226LDM cells cultured in a T75 flask were trypsinized, washed twice with PBS and then lysed by vortexing in BF2 (25 mM Tris/Hepes - pH 8.0, 2 mM EDTA, 0.5% Tween20, 0.5 M NaCl, 1:100 Halt protease inhibitor cocktail). The lysate was incubated on ice for 15 min and then equal volume of BF1 (25 mM Tris/Hepes - pH 8.0, 2 mM EDTA, 0.5% Tween20 and 1:100 Halt protease inhibitor cocktail) was added. For immunoprecipitation, the cell lysate was pre-cleared by incubating 500 μl of the lysate in 50 μl of preblocked Protein A/Sepharose beads for 30 minutes at 4°C on a rotor shaker. The sample was then centrifuged at 200 x g for 1 minute at RT and the pre-cleared supernatant was transferred into a fresh centrifuge tube. 50 μl of the sepharose beads were added to the pre-cleared lysate along with the anti-CTCF antibody (Millipore, 07-729, lot # JBC1903613, pre-screened as described in the previous section) and the samples were incubated overnight at 4°C on a rotating wheel. On the following day, the immune-complexes were recovered by centrifugation at low speed and the supernatant was removed. The pellet was washed three times with immunoprecipitation buffer (BF1+BF2) and each time the beads were collected with centrifugation at low speed. The sepharose was then lysed in SDS-lysis buffer and analysed by SDS-PAGE and western blot analysis as described in previous section.

### 2.4. ChIP-seq

ChIP was performed using the ZymoSpin kit (Zymo Research USA) following the manufacturer’s instructions. In brief, 5 × 10^6^ of 226LDM cells from the control and the treated populations were crosslinked with formaldehyde. The crosslinking was quenched with glycine and the cells were washed twice with PBS with the addition of a protease inhibitor cocktail before pelleting at 1000 g for 1 min at 4°C. The pellet was lysed in Chromatin Shearing Buffer and sonicated using Bioruptor Plus (Diagenode) on high power to obtain fragments of 250-300 bp. ChIP reaction mixes containing sheared chromatin, Chromatin Dilution Buffer, anti-CTCF antibody (Millipore, 07-729, lot # JBC1903613 or no-antibody for negative control) and protease inhibitor cocktail were incubated rotating overnight at 4°C. The next day, ZymoMag Protein A beads were added to the mix and incubated for 1 h at 4°C. The complexes were washed with Washing Buffers I, II and III and then the beads were re-suspended in DNA Elution Buffer. Following de-crosslinking with Proteinase K at 65°C, the ChIP DNA was purified using the ZymoSpin IC columns. The samples were stored at −80°C. The concentration of DNA in the ChIP samples was measured using the NanoDrop 3300 fluorospectrophotometer (Thermo Scientific) along with the Quant-iT™ PicoGreen ds DNA assay kit according to the manufacturer’s instructions. Illumina 50-bp paired-end read sequencing was performed for two biological replicates for each cell state for each antibody, as well as no-antibody Inputs. The sequencing was performed using standard Illumina protocols at the University College London (UCL) Genomics Centre.

### 2.5. RNA extraction

Total RNA from 226LDM cells (three biological replicates from the control and three from the treated population) was extracted using the TRIsure reagent (Bioline) according to the manufacturer’s guidelines. Briefly, cells grown in a T75 flask were washed twice with PBS, then scraped off and pelleted at 300 g for 5 min. Following incubation with TRIsure for 5 min at RT, chloroform was added and the sample was incubated for 15 min at RT. After centrifugation at 9,500 g for 15 min at 4°C, the top aqueous layer was carefully extracted and the genetic material was precipitated with isopropanol for 20 min on ice. After centrifugation (9.500 g / 15 min / 4°C) the pellet was washed twice in 75% ethanol before air-drying the obtained RNA pellet. The RNA was solubilized in sterile water (40-50 μl) and heated for 10 min at 55°C. The pellet was stored at −80°C. The RNA quality was tested using the Agilent Bioanalyzer system; the electropherographs are shown in Supplemental Figure S2. The library preparation and sequencing using the Illumina platforms were performed at the University College London (UCL) Genomics Centre. 50-bp paired-end reads were sequenced for three biological replicates for each of the two cell states resulting in 20-30 million mapped reads per replicate.

### 2.6. ChIP-seq analysis

Reads were aligned to the human hg19 genome with the help of Bowtie [34] allowing only uniquely mapped reads and up to 1 mismatch, resulting in ~72% of total reads being mapped. Around 25-30 million uniquely mapped reads were obtained from each of two replicate experiments, resulting in 50-60 million reads per condition. Mapped reads from two replicate experiments were merged together for each condition before peak calling. Peak calling was performed with MACS 1.4 [35] with default parameters (P=1e-5), using the corresponding Input (no-antibody control) for each experiment. The intersection of genomic intervals were performed using BedTools [36]. Coordinates of CTCF binding sites in MCF-7 determined by the ENCODE consortium were downloaded from the GEO database (GSM822305). Promoter coordinates were obtained from the RefSeq database. The profiles of selected regions and genome-wide aggregate profiles were calculated using NucTools [37] and visualised using OriginPro (Origin Lab) as described previously [37]. Average aggregate occupancy profiles were normalised to 1 at the leftmost end as was done previously [38]. Sequence motif analysis was performed using HOMER [39]. Precise positioning of CTCF binding sites within each category of CTCF peaks was done by scanning for the CTCF motif from JASPAR [40] using RSAT with default parameters [41]. K-means clustering heat maps were generated using NucTools as described previously [37].

### 2.7. RNA-seq analysis

Reads were aligned using Novoalign 3.2 to the reference genome (hg19) and the raw counts were normalized to RPKM values using the Bam2rpkm tool from Galaxy. Differential expression was determined using DeSeq. Genes whose expression change was less than 1.5-fold were included in the “unchanged” gene expression category. Gene Ontology (GO) analysis was performed using DAVID [42], Revigo [43], Cytoscape [44] and Panther [45]. The list of genes that were associated with CTCF binding sites in their vicinity (correspondingly, within +/−10,000bp or +/−1,000bp from TSS as specified in the text) was divided into upregulated/downregulated/no-change based on the RNA-seq data. When a gene was associated with multiple CTCF sites from different classes, it was counted in each of the corresponding classes. The lists of upregulated/downregulated/no change genes associated with CTCF binding sites were intersected with the list of housekeeping genes from [46] in order to determine the enrichment of housekeeping genes in each category.

### 2.8. Data availability

The CTCF and H3K9me3 ChIP-seq as well as RNA-seq data from this study is deposited to the GEO archive (accession number GSE102237).

## 3. RESULTS

### 3.1. 226DM cells treated with hydroxyurea and nocodazole retain CTCF180 and not CTCF130

The 226LDM cell line was chosen as a model to investigate binding patterns of CTCF130 and CTCF180 in the genome, because proliferating 226LDM cells predominantly contain CTCF130, whereas after the treatment with hydroxyurea (HU) and nocodazole (NO) only CTCF180 remains [29]. In addition to our previous work [29], Combined HU/NO treatment is one of standard procedures for mammalian cell cycle synchronization: HU treatment alone arrests cells in G1/S phase, whereas NO-treatment is used to arrest in G2/M phase [7, 47]. HU/NO-treated cells are thus arrested either in G1/S or G2/M; they demonstrate clear morphological changes becoming rounded and suspended in the medium [29, 30]. In addition to our previous works [29, 30], Supplemental Figure S1 (panels A-C) further confirms that CTCF180 can be detected exclusively in normal primary breast tissues, whereas both CTCF180 and CTCF130 are present in MCF7 (cancer) and 226LDM (immortalized) breast cells.

Due to batch-to-batch variations specific screening procedures are required to select the appropriate antibodies that can recognise both CTCF130 and CTCF180 [29]. Such tests were conducted in the current investigation and the antibodies which could recognize both CTCF130 and CTCF180 (Millipore, 07-729, lot # JBC1903613) were selected from the panel of several anti-CTCF antibodies (unsuccessful antibodies are not listed). Using these antibodies we established that HU/NO-treated 226LDM cells lost about 83% of all CTCF signal, and the remaining CTCF is entirely in CTCF180 form (Supplemental Figure S1, panels D-E). As we have previously showed using HU/NO-treated 226LDM cells the CTCF180 form is unequivocally the PARylated CTCF form [29]. Our selected antibodies are able to immunoprecipitate both CTCF forms in untreated 226LDM cells, as shown in Supplemental Figure S3. These antibodies were then used for ChIP-seq analysis of CTCF binding in control (proliferating) and arrested (HU/NO-treated) cells. It is noted that the selection for the required recognition of both CTCF130 and CTCF180 has decreased the overall antibody efficiency in comparison with standard antibody batches used previously by us and others in classical CTCF ChIP-seq (as seen below by the smaller number of detected CTCF sites and weaker ChIP-seq peak shapes). This is the compromise which had to be made in order to study the CTCF130/180 switch.

An additional complication of the following analysis is due to a notable ~83% reduction of CTCF associated with the change in the biological states occurred in treated cells (Supplemental Figure S1, panel D). Importantly, the CTCF mRNA levels in treated cells maintained at 59% in comparison with control cells (Supplemental Table 1). Such moderate variations of CTCF expression are quite common e.g. during cell differentiation, and the associated differences in CTCF binding between the corresponding cell types are not dramatic, meaning that a 41% reduction of CTCF expression by itself would not explain noticeable elimination of DNA-bound CTCF [13, 48]. On the other hand, a reduction of available CTCF would prioritise CTCF binding to stronger DNA sites over weaker sites. It is also worth noting that ~83% reduction of the amount of CTCF proteins does not represent a significant technical challenge for ChIP- and ChIP-Seq experiments and such experiments have been recently successfully conducted in cell lines even with a close to complete CTCF knockout [49].

### 3.2 Analysis of CTCF binding and gene expression profiles in proliferating (control) and arrested (treated) 226LDM cells

The analysis of total transcriptomes of control and treated cells revealed that 2,651 genes were differentially expressed in treated cells (adjusted P-value < 0.05, log2 fold change > 1.5). Among them 1,270 were up-regulated and 1,381 down-regulated. Gene Ontology analysis performed for ranked genes is shown in Supplemental Figures S4 and S5. The changes identified in the transcriptomes were consistent with the two biological states of the cells (proliferating *vs* arrested). Thus, genes involved in cell cycle arrest, differentiation and energy reserve metabolic processes were among up-regulated in treated cells, whereas genes associated with metabolic and cell signaling pathways, ion transport and cell adhesion were down-regulated. In addition, RNA metabolic processes were affected in the latter group of genes.

The analysis of CTCF ChIP-seq revealed that the number of detected CTCF binding sites was considerably higher in control cells (n=9,986) compared to treated cells (n=2,271). The reduction of the number of ChIP-seq peaks in treated cells was consistent with ~83% decrease of CTCF protein content in the nucleus (Supplemental Figure S1E) and ~40% decrease of the ratio of nuclear versus cytoplasmic CTCF content upon the cell cycle arrest (Supplemental Figure S6). The intersection of CTCF binding sites obtained in our experiments with CTCF sites identified in breast cancer cells MCF7 by the ENCODE consortium [50], reveals the overlap of 67% and 19.6% of CTCF sites in control and treated cells, respectively. The high percentage of the overlap in control 226LDM and MCF7 cells confirms the specificity of our ChIP-seq experiment. A lower percentage of the overlapping CTCF sites in treated 226LDM *vs* MCF7 reflect the specific effect of CTCF redistribution upon cell treatment.

We have distinguished three groups of CTCF sites with different binding patterns in control and treated cells, which were termed “common”, “lost” and “gained”. Common sites were bound by CTCF in both cell states. Lost sites were bound by CTCF in control but not in treated cells. Gained sites were only observed in treated cells (Figure 1A). The majority of sites were lost after treatment, and only 257 common sites were retained (Figure 1B).

**Figure 1.**
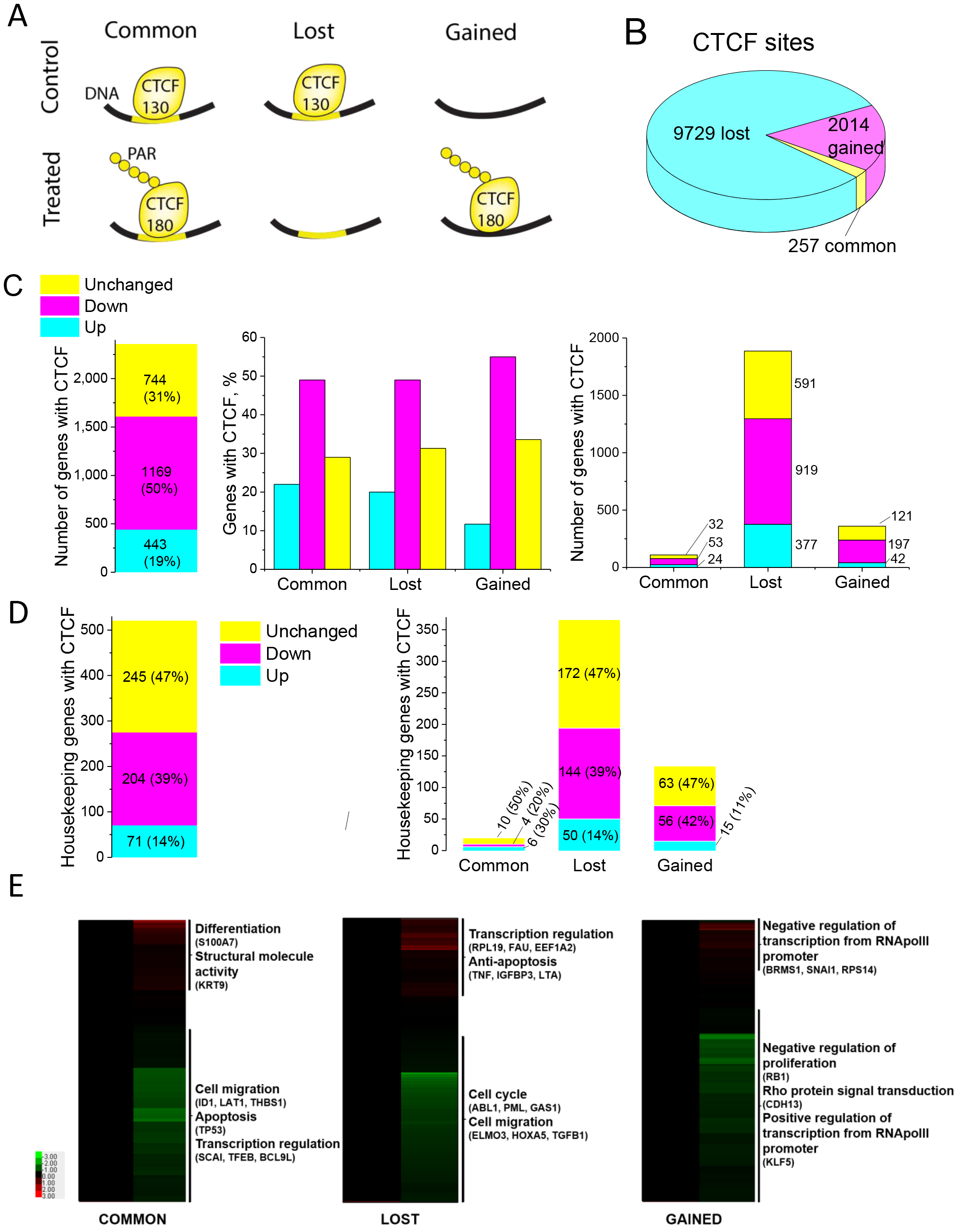
Analysis of CTCF binding and gene expression profiles in control and treated 226LDM cells. (A) Schematic illustration of three groups of CTCF sites detected by ChIP-seq in control and treated cells: common sites are present in both cell states, lost sites are present only in control cells, and gained sites appear only in treated cells. (B) A pie chart showing the numbers of common, lost and gained CTCF sites. (C) Association of gene expression patterns in control and treated cells with the three groups of CTCF binding sites present within +/− 10,000 bp from TSS. Numbers of genes up-regulated, down-regulated and unchanged is shown for all genes near CTCF according to the above criterium (left). Percentages and numbers of genes with different expression patterns for each group of CTCF-binding sites are shown in the middle and right panels, respectively. (D) Association of gene expression patterns in control and treated cells with the three groups of CTCF binding sites present within +/− 10,000 bp from TSS of housekeeping genes. Numbers of genes up-regulated, down-regulated and unchanged is shown for all genes near CTCF (left). Percentages and numbers of genes with different expression patterns for each group of CTCF-binding sites are shown in the right panel. (E) Gene ontology terms enriched for genes containinig CTCF within +/−10,000bp from TSS. Genes are ordered by expression fold change. Red colour corresponds to up-regulation, green colour – down-regulation.

The enrichment of gene ontology terms of genes with promoters containing CTCF is shown in the Supplemental Figure S7. In the common group, up-regulated genes were enriched in developmental processes and down-regulated genes were enriched in response to metal ions. In the lost group, up-regulated genes were enriched in ion binding and homeostasis processes, whereas most of the down-regulated genes were associated with signal transduction and adhesion processes. In the gained group, up-regulated genes were enriched in those involved in the regulation of macromolecular complexes and membrane transporter activities, whereas down-regulated genes were enriched in genes involved in nucleotide binding and biosynthetic processes.

Since the number of CTCF sites within promoter regions in the previous analysis was quite small (e.g. only 35 common CTCF sites at promoters), at the next step we have extended the area of interest to +/−10,000 bp from transcription start site (TSS). CTCF’s action is known to include the formation of chromatin loops between functional regions and this distance is well within the typical range of CTCF action. Gene Ontology analysis of expression of genes containing CTCF within +/− 10,000 bp from TSS showed that, collectively for all three groups, most of these genes were down-regulated upon treatment (1,169 or 49.6%); 443 (18.8%) were up-regulated and 744 (31.6%) unchanged (Figure 1C, left panel). In comparison with all differentially expressed genes, this means that genes associated with CTCF were on average stronger downregulated (Fisher P-value<0.00001). When genes containing CTCF in their vicinity were split according to the status of that CTCF site (common, lost and gained), a similar pattern emerged: the majority of CTCF-associated genes (~50-55%) were down-regulated and ~10-20% were up-regulated (Figure 1C, middle and right panels). Thus, most CTCF-associated genes lost CTCF and decreased their expression upon treatment. At the same time, expression did not change for a large number of genes in these groups (~30-35%). Interestingly, a large proportion of genes CTCF-associated genes are housekeeping according to the classification of Eisenberg and Levanon [46]. Unlike the majority of CTCF-associated genes which where downregulated (Panel 1C), most housekeeping CTCF-associated genes did not change their expression (Figure 1D).

Gene ontology analysis of transcriptional changes revealed genes highly up- or down-regulated in three different groups (Figure 1E). In the common group, highly up-regulated genes were associated with differentiation and down-regulated genes – with cell migration and apoptosis. In the lost, the largest group, highly up-regulated genes were enriched in categories associated with anti-apoptotic processes, whereas most of the highly down-regulated genes were associated with cell cycle and cell migration processes. In the gained group, both highly up-regulated and down-regulated genes were enriched in categories regulating RNA Pol II transcription.

### 3.3. Relationship between CTCF occupancy and gene expression in control and treated cells

Next, we investigated the relationship between the changes in CTCF binding and gene expression. By stratifying all genes containing CTCF within +/−10,000 bp from TSS according to their expression level, we observed that it was more likely to find CTCF in the vicinity of a higher expressed gene in both control and treated cells (Figures 2A and 2B). Furthermore, due to the loss of CTCF near many low-expressed genes upon treatment, this effect is more pronounced in treated cells (Figure 2B). Interestingly, when we stratified genes by their expression fold change upon treatment (Figure 2C), it appeared that there was a clear preference for retained CTCF at common sites to be associated with genes which did not change or changed their expression minimally. Genes considerably up- or down-regulated upon treatment have lost CTCF (see the leftmost and rightmost parts of Figure 2C).

**Figure 2.**
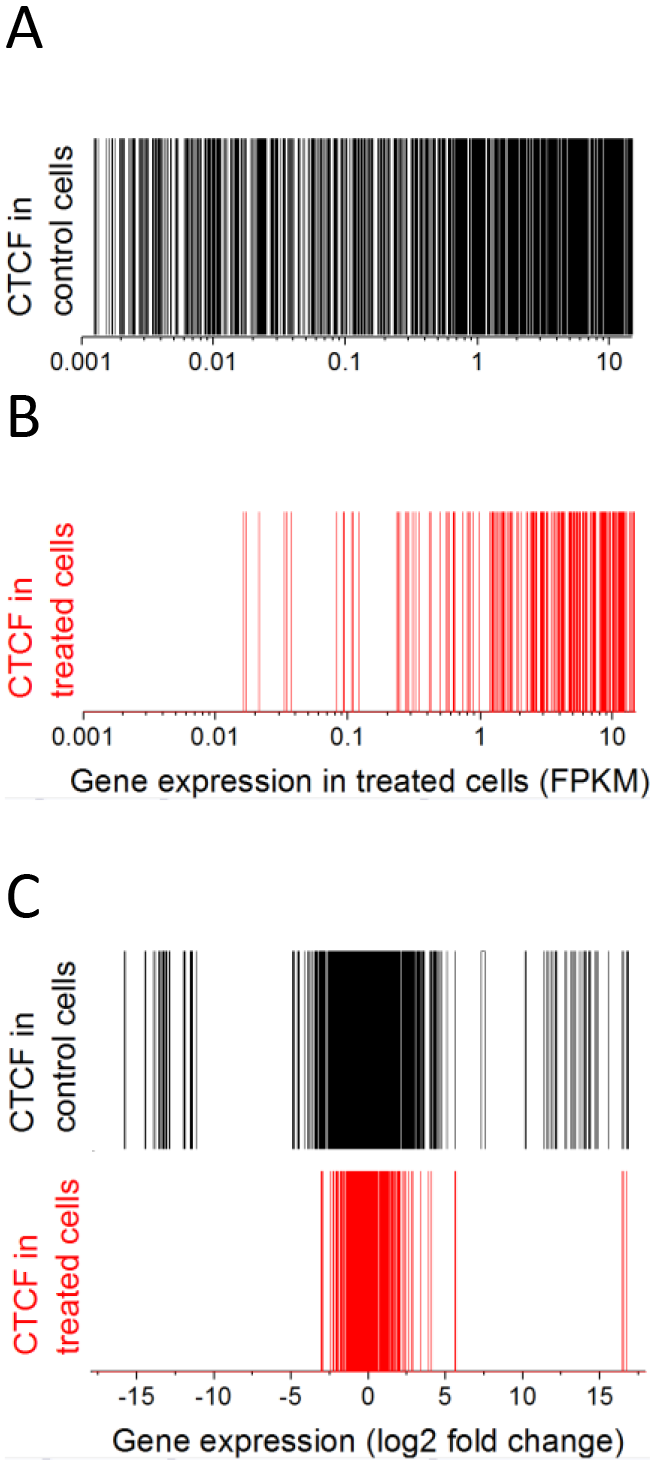
Correlation of CTCF binding within +/−10,000 bp from TSS with gene expression from the corresponding genes. (A) Genes containing CTCF at their promoters in control cells sorted by expression in control cells. (B) Genes containing CTCF at their promoters in treated cells sorted by expression in treated cells. (C) Genes containing CTCF at their promoters in control (black) or in treated cells (red), sorted by expression fold change between treated and control cells. Each vertical bar corresponds to one gene.

We also correlated changes in gene expression with the changes in CTCF occupancies for the three groups of CTCF sites within a more narrow window +/−1,000 bp from TSS, which is representative for transcription factor binding at promoters [51]. In agreement with observations above (Figure 2C), for the group of genes contained common CTCF sites at their promoters the changes in gene expression were relatively small (Figure 3A), while promoters which lost or gained CTCF were associated with a much broader range of gene expression levels (Figure 3, panels B and C, respectively). No correlation was observed between CTCF occupancy and gene expression in the common and lost groups, although small positive correlation (r=0.15) was seen in the gained group.

**Figure 3.**
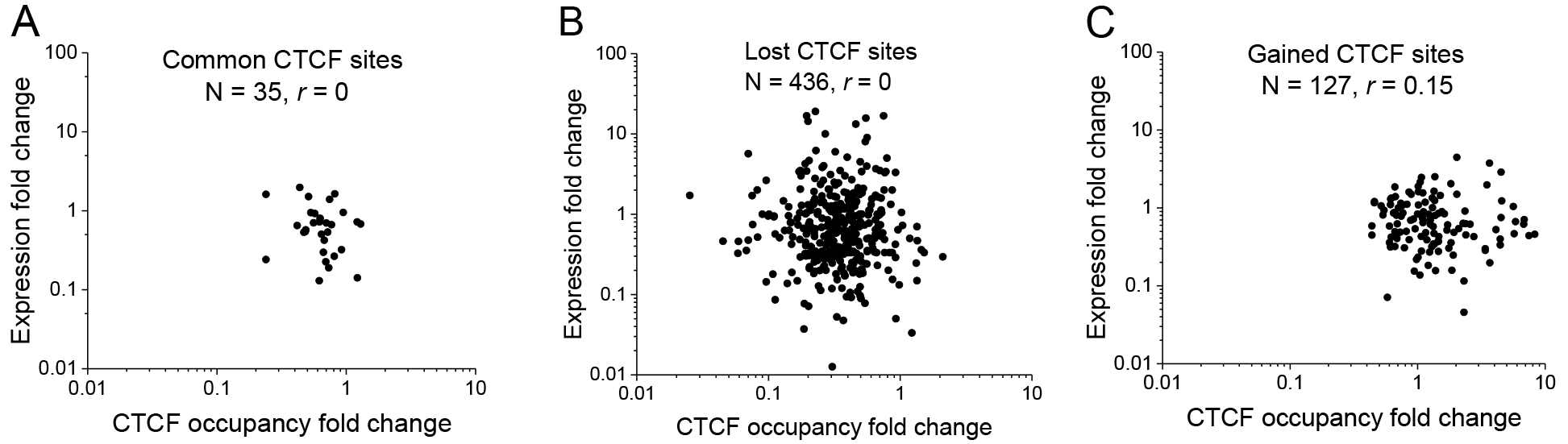
Correlation of gene expression fold change and CTCF occupancy change at the corresponding promoters (+/−1,000 bp from TSS) for the common (A), lost (B) and gained (C) CTCF sites. Promoters with common and lost CTCF sites did not show statistically significant correlation of the change of gene expression with CTCF occupancy change upon cell treatment.

### 3.4. Common and lost, but not gained CTCF sites contain classical CTCF binding motifs

CTCF employs a combination of its eleven Zinc fingers to bind to diverse DNA sequences with a consensus ~19 bp motif [52]. Most of CTCF sites contain the classical consensus sequence, but CTCF sites with different consensus motifs and those which do not match any consensus motifs have been also reported previously [52–55]. To identify PARylation specific features we calculated the nucleotide frequencies as a function of distance from the summit of CTCF ChIP-seq peak for the subsets of the sites from the common, lost and gained groups. As shown in Figure 4, CTCF sites in the common and lost but not the gained groups contain classical CTCF recognition motif, enriched with the guanine and cytosine residues at the summit, although this pattern is more pronounced for the common sites. Interestingly, the nucleotide distribution in the 3’ and 5’ flanking regions of the motifs significantly differs between these groups, demonstrating higher GC content in the common group. This is in line with our previous observation that common but not lost/gained sites were enriched inside CpG islands for the system of mouse embryonic stem differentiation [38].

**Figure 4.**
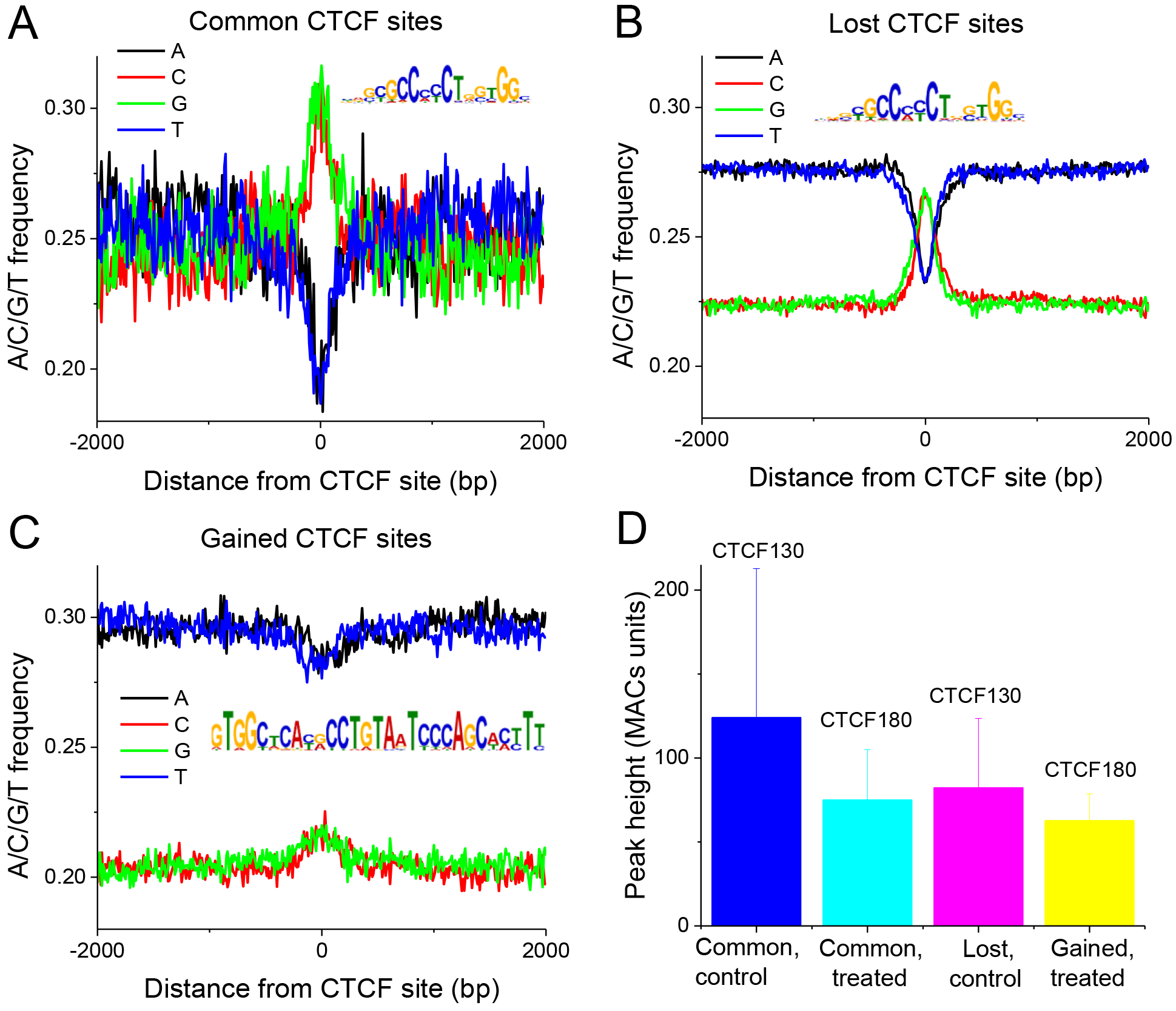
Nucleotide frequencies as a function of distance from the summit of CTCF ChIP-seq peak for the subsets of common (A), lost (B) and gained sites (C). The consensus motifs for common, lost and gained CTCF sites are shown in the inserts in the corresponding panels. Most common CTCF sites contain classical CTCF recognition motif; part of lost CTCF site also contain the classical CTCF recognition motif; gained CTCF sites do not contain CTCF binding motif, suggesting nonspecific binding to open chromatin regions defined by other TFs. D) The strength of CTCF binding reflected by the heights of ChIP-seq peaks for different classes of CTCF sites.

We have also assessed the strength of CTCF binding for different classes of CTCF sites calculated by the heights of the ChIP-seq peaks (Figure 4D). The strongest binding was observed in the control cells in the common group, which on average almost did not change after treatment. The initial CTCF signal in the lost group was smaller than in the common sites before treatment, and it significantly decreased after treatment. The lowest signal was in the gained group which most likely reflected the nature of CTCF180 binding (very weak or DNA-independent).

From these analyses we conclude that common and lost sites are characterised by the presence of the CTCF consensus motif. The strongest CTCF binding is observed for common sites, whereas it is weaker for the lost sites. Gained sites have no classical CTCF motif and their CTCF binding is the lowest as expected. Thus, gained sites may represent nonspecific interactions of CTCF with chromatin or regions where CTCF is indirectly bound to DNA.

### 3.5 Loss of CTCF binding upon cell treatment is associated with nucleosome repositioning

The importance of CTCF in the regulation of the nucleosome distribution has been widely recognized [38]. CTCF binding also often demarcates distinct chromatin states and protects DNA from methylation [4, 8, 56]. Previous studies showed that CTCF regularly positons several nucleosomes in its vicinity, resulting a characteristic oscillatory pattern of nucleosome occupancy around CTCF [38, 57–59]; this oscillatory nucleosome pattern disappears when CTCF binding is lost and nucleosome depletion at the center of CTCF site is then replaced by a strongly positioned nucleosome [58]. Thus, it was interesting whether the CTCF130/180 switch investigated here was associated with some changes in nucleosome positioning. In our first analysis, we considered the DNA protection from shearing in the no-antibody Input sample as an indicator of nucleosome occupancy, as was done in several previous studies [60, 61]. It is important to note that while a number of factors other than nucleosome occupancy can also affect the Input read coverage distribution, the irregularities in the Input coverage landscape observed at *mono-nucleosome scale* mostly represent the nucleosome resistance to the sonication [62]. Essentially, at the mono-nucleosome scale ChIP-seq Input reads density reflects the nucleosome occupancy in a very similar way as MNase-seq; the difference from MNase-seq is only the resolution of nucleosome positioning. Similar to MNase-seq and many other sequencing techniques, ChIP-seq Input also has sequence-dependent artifacts of which we are aware [63–65]. In order to increase the resolution of nucleosome occupancy obtained from ChIP-seq Input one can plot the density of “plus tags” – the start coordinates of the reads. Furthermore, the resolution of positions of CTCF binding sites around which nucleosome positioning is considered can be improved by substituting CTCF ChIP-seq peaks by exact locations of CTCF sites within CTCF ChIP-seq peaks found by scanning for the CTCF motif.

Supplementary Figure S8A shows the average profile of the plus tag density around CTCF sites within CTCF ChIP-seq peaks lost in treated cells, defined with single-base pair resolution by CTCF motifs. This figure unequivocally shows that the read density reflects the nucleosome occupancy: in control cells the plus tag density around CTCF sites shows characteristic oscillations as in standard MNase-seq experiment, while in treated cells CTCF sites that lost CTCF binding also lost the nucleosome oscillation pattern. Furthermore, nucleosome depletion at CTCF site observed in control cells is replaced by a nucleosome occupancy peak in treated cells. The same effect is observed in the total read density when DNA reads were extended by the average read length, although in this case the effect is blurred (Supplementary Figure S8B).

We then performed similar calculations for nucleosome occupancy around common/lost/gained CTCF peaks without refining CTCF sites to CTCF motifs. Figure 5 shows that CTCF130 bound regions in the common control group are associated with smaller average nucleosome occupancy than the same regions in treated cells. This also correlated with the reduced strength of CTCF180 binding at these sites after treatment (Figure 4D). In the lost group, average nucleosome occupancy increased after treatment and release of CTCF180 (Figure 5B), whereas in the gained group nucleosome occupancy at CTCF180 binding sites did not change following CTCF recruitment (Figure 5C). Taken together, these findings indicate that CTCF binding and nucleosome occupancy at its binding site are anti-correlated. This data is consistent with a number of previous reports on the competition of CTCF with nucleosomes *in vivo* [38, 59] and with the observation that that the regions including CTCF site in human cells contain an intrinsic nucleosome positioning signal for a single nucleosome centered at the CTCT site [66].

**Figure 5.**
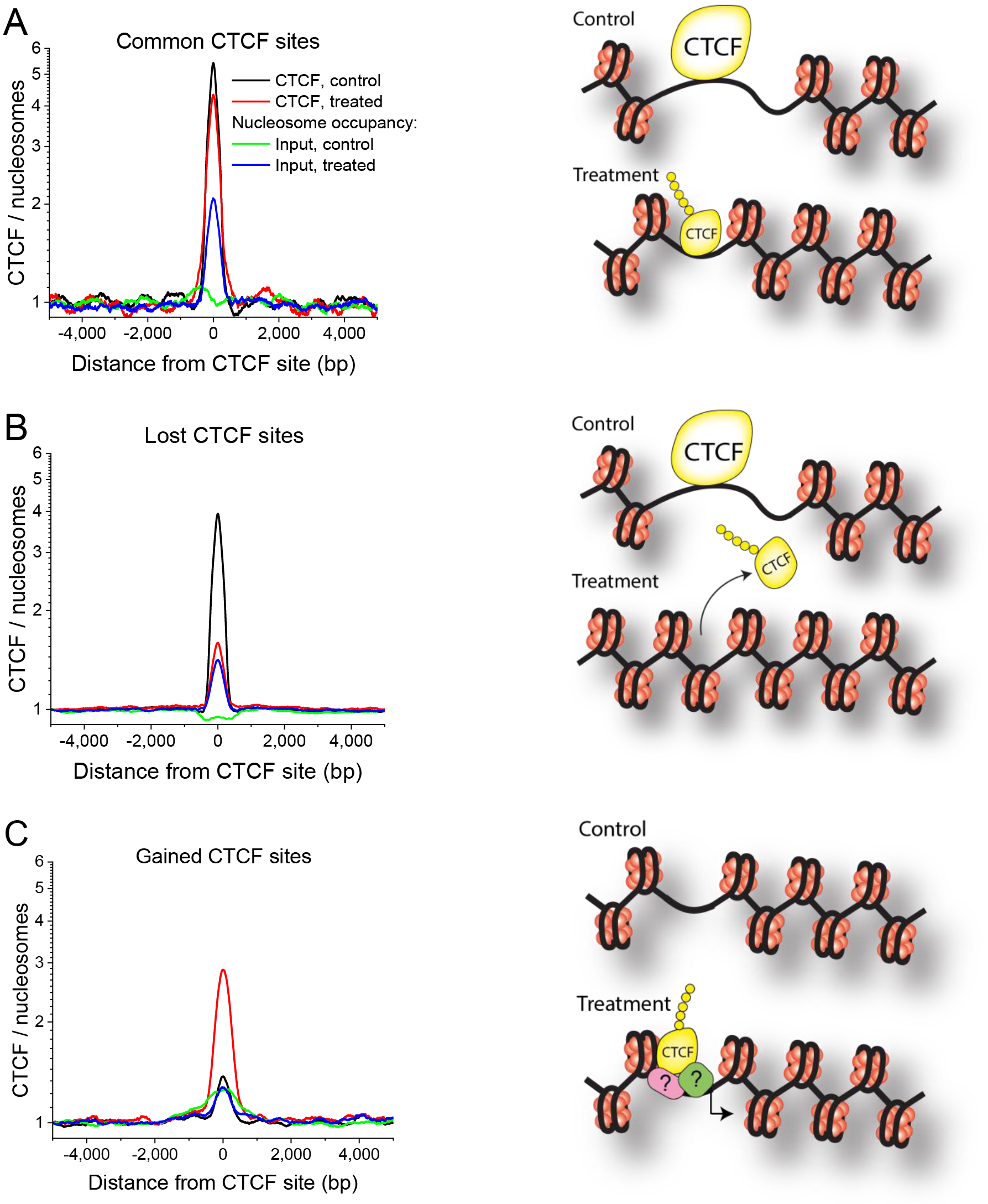
Average profiles of CTCF and nucleosome occupancy at common (A), lost (B) and gained (C) CTCF sites. Black – CTCF in control cells, red – CTCF in treated cells, green – nucleosome occupancy in control cells, blue – nucleosome occupancy in treated cells.

Nucleosome redistributions reported in Figure 5 can be associated with different post-translational histone modifications. As a test case, we have performed ChIP-seq using anti-H3K9me3 antibody in untreated and treated cells. H3K9me3 has been selected because in our previous work it was shown to be associated with higher density of mapped reads and higher nucleosome density [58]. As shown in Figure 6 (panels A and B), a significant H3K9me3 redistribution occurs around common and lost CTCF sites. Interestingly, no such rearrangements around gained CTCF sites were observed, which suggests that gained sites may be non-specific/non-functional.

**Figure 6.**
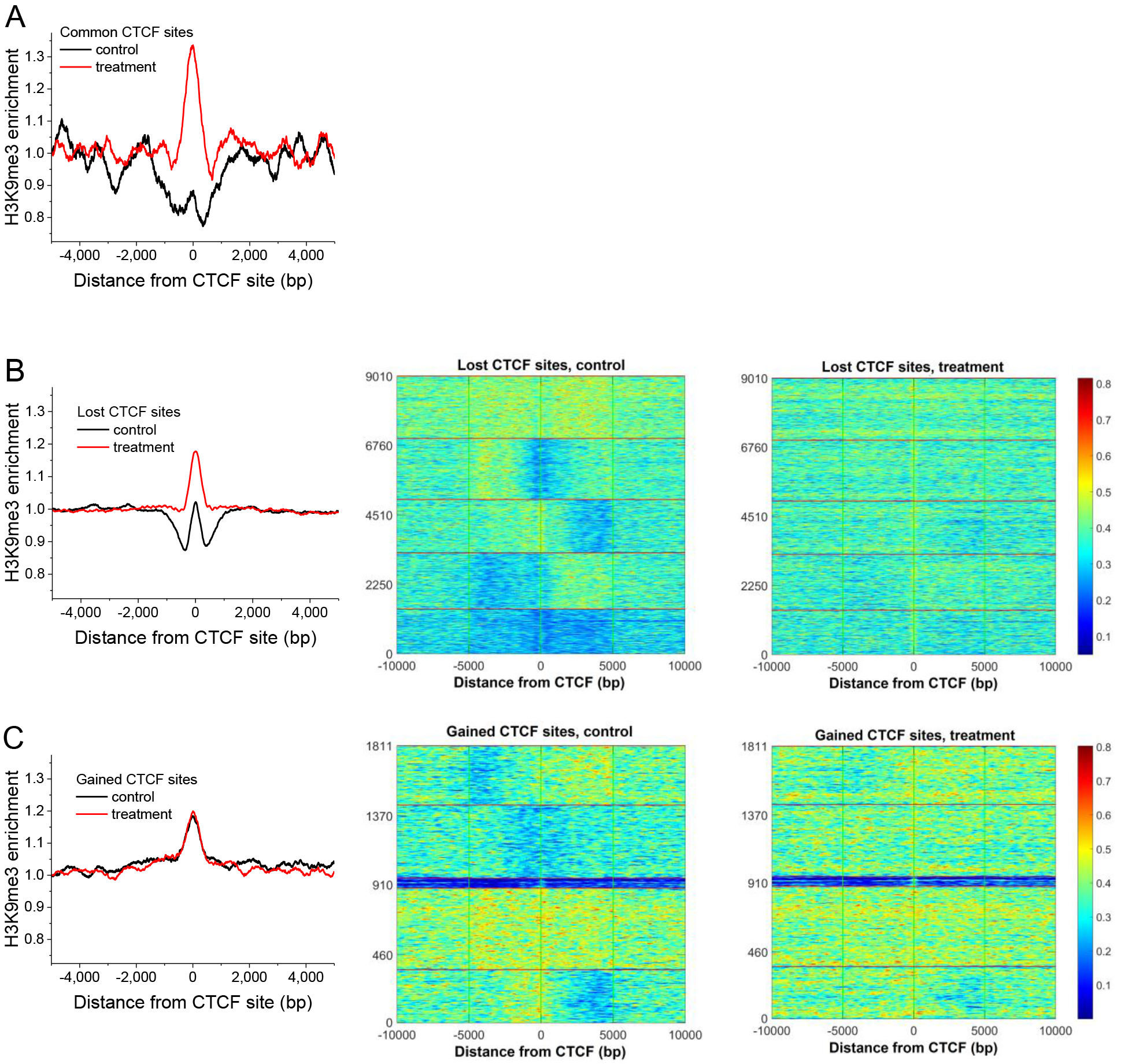
Average profiles and k-means clustering heat maps of H3K9me3 enrichment at common (A), lost (B) and gained (C) CTCF sites. Black -control cells, red – treated cells. In the case of common sites the number of sites was not enough for k-means clustering and the corresponding heat maps are not shown. In the case of lost an dgained sites, k-means clustering was performed for the control condition. The region ordering heat maps for the treated cells correspond to the same order as in the corresponding heatmaps for the control condition.

Since PARylation can change physical CTCF interactions with chromatin proteins, we have also looked at the chromatin density profiles based on the Input read density near individual CTCF sites (not at the CTCF binding site itself, but rather in the immediate vicinity). Examples of specific gene promoter regions where CTCF-associated chromatin rearrangements take place following treatment, together with changes in gene expression patterns, are given in Supplemental Figure S9. We noted that in many cases CTCF binding in control cells was associated with sharp Input peaks in the physical proximity to CTCF, which disappear in treated cells. In addition, in some cases the Input and CTCF peaks are situated at different ends of the gene (e.g. an Input peak at the transcription start site (TSS), and CTCF at the transcription end site (TES) in the case of E2F4 (Supplemental Figure S9). The latter suggests possible TSS-TES bridging by CTCF in control cells, which disappears after treatment. The effect of CTCF-dependent chromatin reorganization was observed for all three groups of sites (common, lost and gained), and was not correlated with changes of gene expression (gene expression could go either up or down following treatment). The fact that CTCF-dependent chromatin peaks were next to CTCF but did not coincide with it provides an argument that the chromatin peak is not formed by CTCF itself. Cell treatment-dependent depletion of Input peaks near CTCF sites at the PARP3 and TP53 promoters was also confirmed by ChIP experiments using the DNA primers for the regions at the summits of the corresponding ChIP-seq Input peaks (panels D and E in Supplemental Figure S9). We also analyzed profiles of H3K9me3 chromatin marks in the same promoters as in Supplemental Figure S9. As shown in Supplemental Figure S10, the strength of H3K9me3 signal increases around the regions near CTCF sites which lost chromatin peaks. This suggests that perhaps the lost sharp Input peaks represent specific chromatin complexes other than nucleosomes.

## 4. DISCUSSION

This study aimed to analyse the effect of the CTCF130/180 switch on chromatin structure and gene expression. In the absence of a specific anti-CTCF180 antibody, it was rational to use the 226LDM cell line in which a switch from CTCF130 to CTCF180 can be induced and validated using anti-CTCF antibodies recognizing both CTCF130 and CTCF180. Following the optimization of hydroxyurea and nocodazole concentrations [30], it was possible to obtain viable treated cells with CTCF180, whereas CTCF130 was predominantly present in proliferating control cells. Following the cell cycle arrest we observed ~83% decrease of CTCF protein content in the nucleus (Supplemental Figure S1E) and ~40% decrease of the ratio of nuclear versus cytoplasmic CTCF (Supplemental Figure S6). Such a pattern of CTCF distribution was previously reported in normal breast tissues where only CTCF180 is detected. Interestingly, a transition from CTCF180 to CTCF130 took place in primary cultures generated from normal cells from breast tissues indicating the labile nature of this modification [29].

The generation of CTCF180 in response to the drugs can be explained by the initiation of checkpoint signalling cascades, leading to activation of PARP enzymes and subsequent PARylation of CTCF. Indeed, the nocodazole- [67] and hydroxyurea-induced [68] cell cycle arrests have been linked to the activation of the PARP-signalling pathways [69, 70]. Global changes in gene expression profiles were consistent with the changes in the biological states of the cells, revealing up-regulation of genes involved in cell cycle arrest, development, differentiation and energy reserve metabolic processes and down-regulation of genes associated with metabolic and cell signaling pathways, ion transport and cell adhesion (Supplemental Figures S4 and S5).

Our ChIP-seq analysis confirmed for the first time that CTCF180 has well-defined genomics targets, paving the way for further research into the specifics of this binding in different conditions, cell lines or tissues. The number of CTCF180 sites detected in treated cells was found to be much smaller than in control cells (n=2,271 vs n=9,986, respectively), which is explained by the reduction of the total CTCF concentration in chromatin (Supplemental Figure S6). The remaining smaller number of common CTCF sites in treated cells may implicate that they are involved in the organization of 3D chromatin structure, and thus have higher affinity and are surrounded by other cooperatively interacting proteins. Moreover, the protein composition of CTCF-interacting complexes is likely to be different because of the particular nature of CTCF180. These aspects will need to be explored in the future, especially for the primary tissues where CTCF180 is naturally very abundant (and in some tissues, such as breast, it is the only form).

This study provides new insights in DNA-binding and gene regulatory properties of CTCF180 summarized in Figure 7. Our results suggest that common and lost sites contain the classical CTCF motif, although the former are more GC-rich at the summit and in the background around the motif, whereas the latter are embedded into more AT-rich sequences. A similar effect of common CTCF sites residing in more GC-rich and CpG-rich areas has been previously noticed in our study of mouse embryonic stem cell differentiation [38] and it seems to be a general effect. The effects of flanking DNA sequences regions may be also linked to the strength of CTCF binding, which is highest for the common sites (Figure 4D). The fact that no CTCF binding motif was observed in the gained group suggests that CTCF-DNA interaction at these sites is non-specific or CTCF180 interacts with these regions in a DNA-independent manner, directly or through recruitment by other proteins.

**Figure 7.**
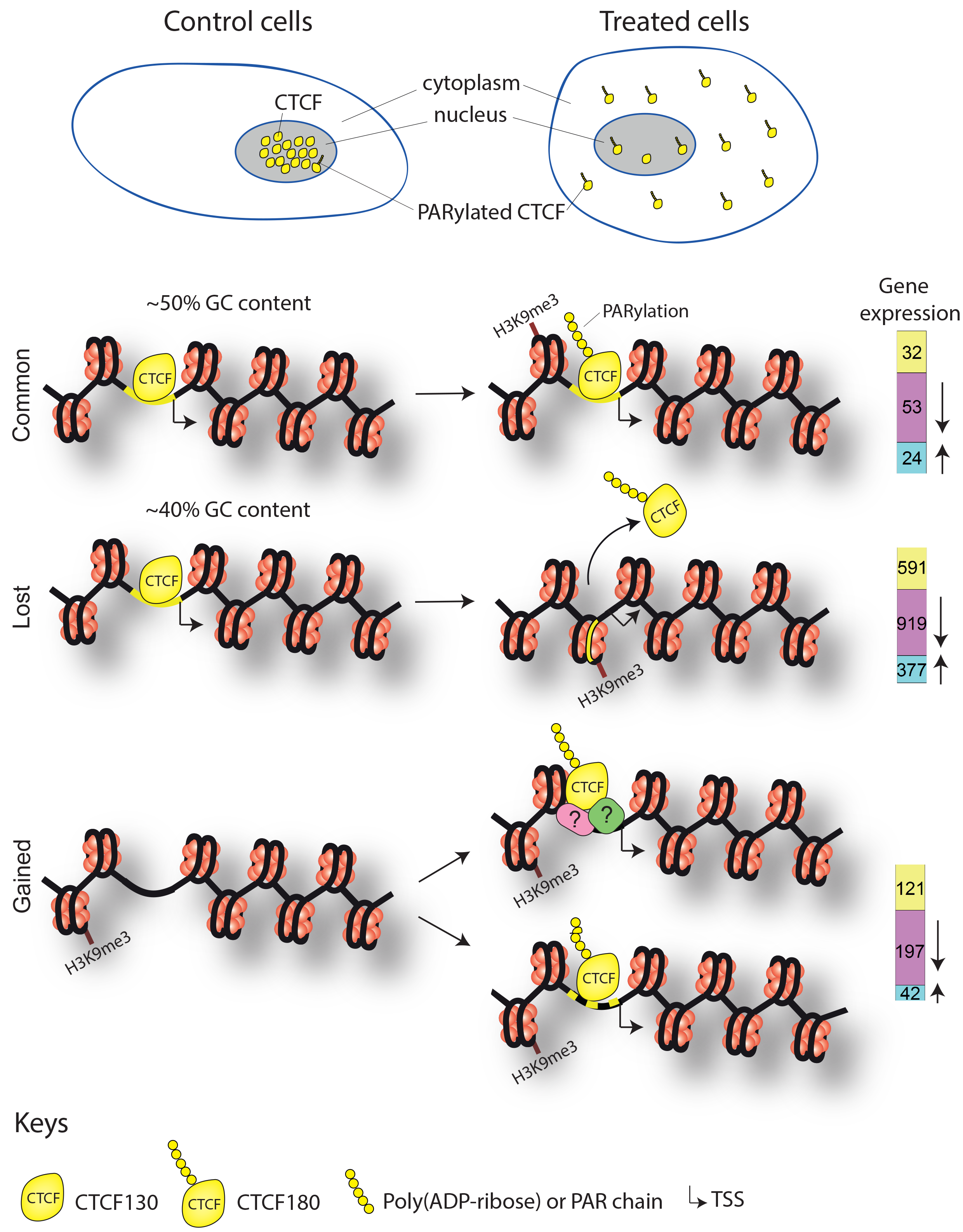
A schematic model illustrating the events observed in control and treated 226LDM cells in which transition from CTCF130 to CTCF180 takes place. Following treatment, cells change morphologically from adherent and flat to suspended and rounded. PARylated CTCF180 in treated cells is largely redistributed from the cell nucleus into cytoplasm (depicted on top of the Figure). More GC-rich stronger common sites retain CTCF180 (with smaller strengths). CTCF180 is evacuated from weaker (lost) sites. Gained sites characterised by the absence of the CTCF motif acquire CTCF180 after treatment possibly due to interaction with additional proteins or may be just false positives. Nucleosome occupancy associated with the higher levels of the H3K9me3 is increased in the regions overlapping with CTCF sites in common and lost groups. Molecular changes within regions containing these CTCF sites result in alterations in gene expression patterns.

The change in nucleosome occupancy resulting from loss of CTCF observed here is similar to the effect of CTCF/nucleosome competition that was previously reported [48, 71, 72] and it could be one of the mechanisms explaining CTCF redistribution. Additional mechanisms can be through treatment-induced changes of chromatin modifications. At least one of histone modifications redistributing at common and lost (but not gained) CTCF binding sites was found to be a heterochromatin mark H3K9me3. The latter is in agreement with down-regulation of the majority of the genes within these regions (Figure 1). On the other hand, many up-regulated genes were also associated with CTCF, implying the involvement of additional regulatory factors/mechanisms [49, 71, 73]. The complexity of this regulation is further illustrated by the observations that changes of CTCF binding at promoters were associated with chromatin rearrangements in the regions adjacent to CTCF binding sites (Supplemental Figures S9 and S10). It did not escape our attention that chromatin rearrangement in the vicinity of PARylated CTCF sites may represent a general effect, but exploring its nature would require additional extensive experiments which are beyond the scope of the current work. Interestingly, the connection of chromatin PARylation to its decompaction has been noted recently, which might be related to the effect reported here [74]. However, one should be also cautious to not over-interpret individual ChIP-seq Input peaks due to possible artifacts [60, 75–77].

The majority of CTCF-associated genes experienced both CTCF loss and gene expression downregulation (Figure 1). Unchanged expression of a significant number of genes that lost and gained CTCF near their TSS indicates that the regulation of some genes does not depend on CTCF binding. This effect is particularly pronounced the case for the common group in which it may be important to sustain the optimal level of expression needed for survival of cells in different functional states (Figures 1D and 3).

The importance of CTCF modification in the biological processes is supported by changes in expression profiles of genes associated with CTCF (Figures 1E and 7, far right). These changes involve down-regulation if genes involved in cell cycle and cell migration, and up-regulation of genes involved in differentiation thereby adequately reflecting the biological situation, i.e. transition from proliferating to arrested cells. Furthermore, some of the affected genes appear to be characteristic for particular groups of CTCF sites. For example, genes responsible for cell cycle regulation are down-regulated in the group of genes where CTCF is lost. It is tempting to speculate that such preference may be due to the change of behaviour of PARylated CTCF at the particular type of CTCF sites.

It should be noted that in this report we investigate local effects of CTCF PARylation on its DNA binding properties and, subsequently, changes in adjacent chromatin regions and associated gene expression. It was beyond our scope to consider in this experimental model 3D effects of CTCF rearrangements on higher order chromatin structures [12, 49, 71, 73, 78–80], which may be a subject of the follow up work.

Interestingly, the effect of CTCF PARylation observed in treated cells was not a direct consequence of increased expression of PARP genes. Indeed, PARP-family genes were either downregulated upon treatment (PARP1: fold-change 0.3, adjusted P=0.01; PARP16: fold-change 0.3, adjusted P=0.02) or did not change their expression significantly (PARP2, PARP3, PARP4, PARP6, PARP8, PARP9, PARP10, PARP11, PARP12, PARP14, PARP15). This is consistent with PARP1 downregulation in G1 arrested cells reported recently [81]. In addition, enzyme PARG that is responsible for de-PARylation was only insignificantly downregulated in treated cells (fold change 0.45, adjusted P=0.38). Therefore, one can speculate that the change of CTCF PARylation is due to changed stability or activity of (de)PARylation enzymes rather than their expression levels. In addition, the lack of PARP1 upregulation suggests that the DNA damage response pathways are not among the major determinants of CTCF relocation. Note also that the promoter of one of the main DNA damage response players, p53, is one of the few genes marked by the common CTCF sites.

The effect of CTCF PARylation studied here may be also considered in the general context of posttranslational CTCF modifications (similar to the language of histone modifications), which may deserve a new systematic study due to the particular importance of CTCF in cell functioning. For example, another CTCF modification, phosphorylation, is abundant during mitosis, and has been also reported to affect CTCF binding affinity to chromatin [82]. CTCF is believed to be retained during mitosis at some but not all sites [83–85], an important subject related to our system, which is still not entirely understood. Furthermore, mitotic bookmarking in general is an active area of research [86]. Another example posing an unresolved puzzle is the apparent disappearance of CTCF during S phase reported for mouse embryonic stem cells [87], which may be explained by some CTCF posttranslational modification that makes it un-detectable with standard antibodies. Our results may help elucidate some of the controversies in the field, which at least in part may be attributed to CTCF changes that make them “invisible” to some antibodies.

Finally, this study issues an important cautionary note concerning the design and interpretation of any experiments using cells and tissues where CTCF180 may be present and can go undetected since not all antibodies can recognize this form of CTCF. The 226LDM cells as a model for the switch from CTCF130 to CTCF180 provided us with a unique opportunity to develop an experimental framework to study CTCF180. This approach can be used to investigate the role of CTCF180 in cell lines and tissues, normal and tumour, where either both forms or exclusively CTCF180 are present. The screening of the existing antibodies for their ability to recognise either both forms of CTCF or CTCF130 only will be necessary to enable to subtract the targets recognised by CTCF130 from the combined CTCF130/CTCF180.

## 5. ACKNOWLEDGEMENTS

We thank Parmjit Jat and Michael O’Hare for 226LDM cells and Mike Hubank and colleagues from the University College London (UCL) Genomics Centre for sequencing and initial bioinformatics analysis of the data. We also thank Adele Angel for excellent technical assistance, Hulkar Mamayusupova for advice on quantification of immuniofluorescent images, and Theodoros Giakoumis and Yemane Tedros for their help with ChIP sample preparations. We thank members of Vladimir Teif’s, Elena Klenova’s and Greg Brooke’s laboratories for many helpful discussions.

## Author contributions

F.D., I.P., V.T. and E.K. conceived the project, I.P., F.D. and E.K. performed experiments. I.P., I.C., C.C., V.T. and E.K. analysed the data, E.K., I.P. and V.T. wrote the manuscript.

## Supporting Materials

### Cell cycle arrest treatment with hydroxyurea and nocodazole

226LDM cells approximately 60-70% confluent were stripped off their spent medium and fresh culture medium containing 100 mM hydroxyurea was added to the flask. The treatment was administered as described by Docquier et al [1]. The cells were incubated for 24 hrs at 37°C and CO_2_. After the end of the incubation time, the cells were incubated in fresh complete medium for 1hr at 37°C and CO_2_. After 1 hr the complete medium was aspirated off and fresh medium supplemented with 500 ng/ml nocodazole was added. After 24 hrs the detached cells were harvested and prepared for western blotting and chromatin immunoprecipitation. Quantification of the bends in western blots was performed using ImageJ [2] following the standard instructions provided by the developers.

### Protein Immunoprecipitation

226LDM cells were trypsinised and then lysed by vortexing in BF2 (25 mM Tris/Hepes - pH 8.0, 2 mM EDTA, 0.5% Tween20, 0.5 M NaCl, 1in100 Halt Protease Inhibitors). The lysate was incubated on ice for 15 min and then equal volume of BF1 (25 mM Tris/Hepes - pH 8.0, 2 mM EDTA, 0.5% Tween20, 1in100 Halt Protease Inhibitors) was added. The cell lysate was pre-cleared by incubating 500 μl of the lysate in 50 μl of pre-blocked Protein A/Sepharose beads for 30 minutes at 4°C on a rotor shaker. The sample was then centrifuged at 200 g for 1 minute at RT and the pre-cleared supernatant was transferred into a fresh centrifuge tube. 50 μl of the sepharose beads were added to the pre-cleared lysate along with the antibody and the samples were incubated overnight at 4°C on a rotating wheel. On the following day, the immune-complexes were recovered by centrifugation at low speed for 1 min and the supernatant was removed. The pellet was washed three times with immunoprecipitation buffer (BF1+BF2) and each time the beads were collected with centrifugation at low speed for 1 minute. The sepharose was then lysed in SDS-lysis buffer and analysed by SDS-PAGE and western blot analysis.

### Immunofluorescence (IF) staining on fixed cells

In immunofluorescence staining, which is based on the same principle as immunocytochemistry staining, an antibody is used to detect a specific protein. This antibody is appropriately tagged with a visible dye such as a fluorescent dye [3]. The IF staining was performed on adherent cells grown on cover slips. The cells were fixed with addition of 4% paraformaldehyde (PFA). After three washes with PBS/glycine (0.1 M), they were incubated for 15 min in boiling citrate buffer (10 mM citric acid, pH 6.0). After incubation for 20 min with permeabilization buffer (0.25% Triton / PBS) the cells were washed thrice with 1 × PBS. The coverslips were placed in petri dishes, circled with hydrophobic marker on the slides and 90 μl of blocking buffer (0.05% Tween, 2% serum, 1% BSA / 1 × PBS) was added. A moist towel was put in the dish to keep the environment humid and the dish was covered and left on slow rocking in 4°C overnight. A 2 h long incubation with primary antibodies in buffer (0.05% Tween, 1% BSA, in 1 × PBS) was followed by three PBS washes. Incubation continued in the dark (covered with foil) with secondary antibodies conjugated with fluorescent dyes (e.g. FITC, TRITC) for 1hr. After 3 more washes with 1 × PBS, the coverslips were left to dry and then they were mounted to microscope slides with DAPI (4’,6-diamidino-2-phenylindole, dilactate) mounting medium.

### Microscopy

Immunofluorescence staining was observed under the Nikon Ti-Eclipse wide-field microscope which was used to capture the staining images. The visualisation tool used to view the images was Fusion FX viewer from Nikon.

### Image quantification

Quantification of fluorescence intensity was performed on immunofluorescence staining images that were acquired with the same microscope settings using Fiji/ImageJ [2]. Briefly, the integrated pixel intensity (mean intensity per area) of the whole cell and the nucleus were measured for control and treated cells. The mean intensity of areas between cells was also measured to serve as background. For three different cells (n=3) from each condition, we calculated and compared the average mean intensity of control and treated cells in the whole cell and nucleus. After subtracting the background intensity in each case, the ratios of the resulting intensities were shown in the form of the box plot.

### qPCR analysis

Real-time PCR was performed using ChIP samples to determine the disappearance of the peaks of chromatin density near CTCF binding sites illustrated in Supplementary Figure S9. The summits of the chromatin density peaks in control cells were determined from the green line Figure S9A for promoters of PARP3 and TP53. The following primer sequences were derived to detect the chromatin peaks for PARP3 and TP53:

PARP3 – FORWARD: TCAGAAGCGCCATGCTCA; REVERSE: AACAGCGGCTGCTCGTAAG;
TP53 – FORWARD: TATTCTCCGCCTGCATTTCT; REVERSE: TTCAAAGAAGGGGAGGGATT;

Each sample was amplified using SYBR green (SensiFAST SYBR no-ROX kit, Bioline) with the Lightcycler 96 instrument (ROCHE, CH). The manufacturer’s instructions were followed regarding the reaction and cycling conditions. Briefly, 1 μl of ChIP DNA and 0.2 μM of primers were used to make up a 10 μl reaction. qPCR was performed separately for Input and for the no-antibody control. The no-antibody control did not produce qPCR signal and was assigned Ct value 40 as per standard instructions. qPCR intensity (fold-change) in the Input was compared between control and treated cells using the ΔΔCt method [4]. Following the ΔΔCt quantification, the chromatin density at the ChIP-seq peak summits near CTCF in control cells was assigned value one, and the density in treated cells was defined as a ratio to that in control cells.

**Supplementary Figure S1.**
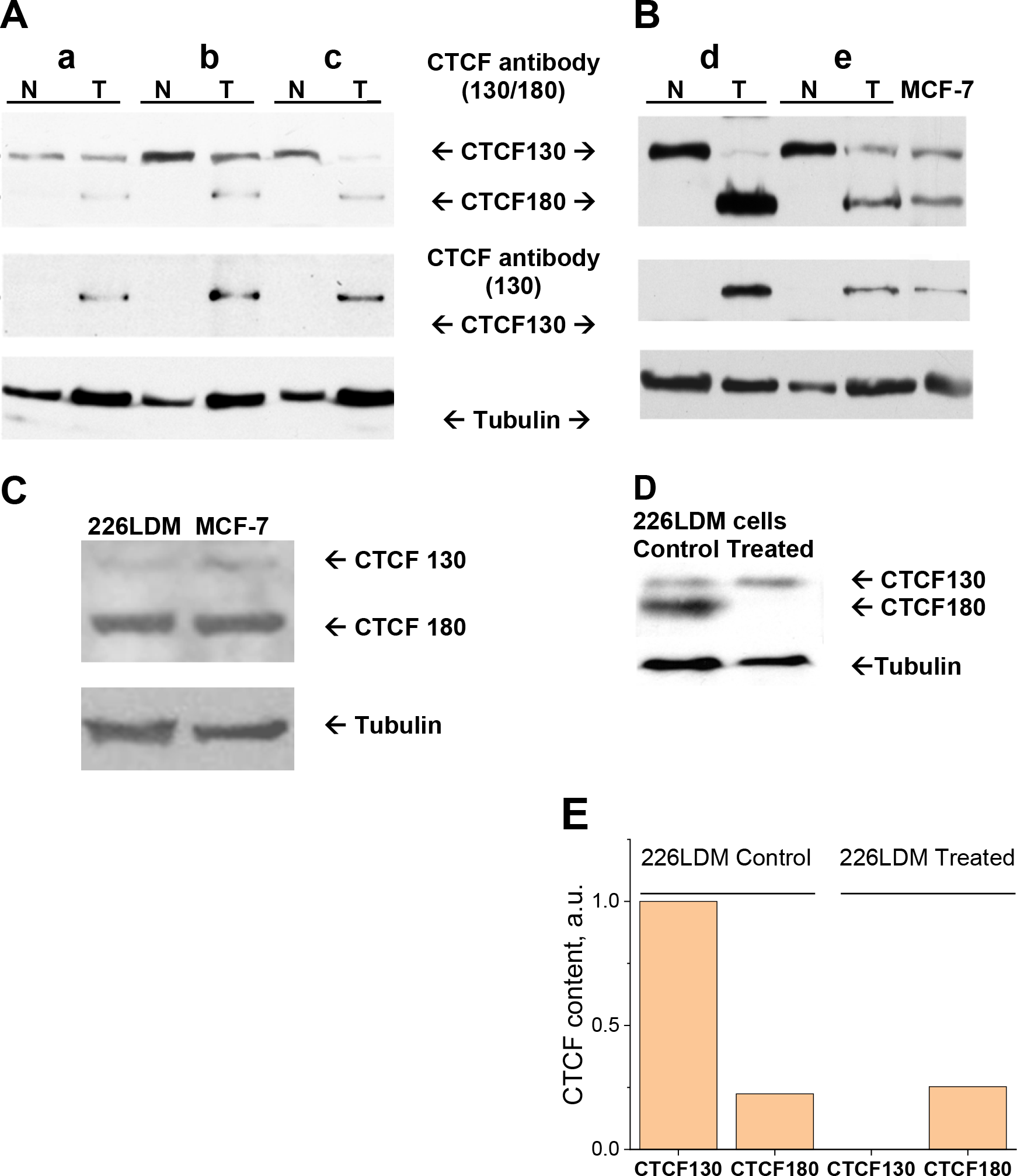
Western Blot analyses of primary breast tissues and breast cell lines demonstrating the presence of two forms, CTF130 and CTCF180, with the antibodies prescreened as previously described [1]. A, B and C: CTCF migrates as 180 kDa protein in normal breast tissues, CTCF-130 appears in tumour breast tissues, and both forms of CTCF are present in MCF-7 breast cancer cells and 226LDM immortalized breast cells. **A.** Western blot analysis of three independent paired samples of normal and tumour tissues (“a”, “b” and “c”). Tissue lysates (50 μg of the total protein) prepared as previously described [5] were resolved by SDS-PAGE, blotted and probed with the pre-screened anti-CTCF polyclonal antibody recognising CTCF180 and CTCF130 (“CTCF 130/180”, upper panel), stripped and re-probed with the anti-CTCF antibody recognising only CTCF130 (“CTCF130”, middle panel), re-stripped and re-probed with the anti-tubulin antibody (“Tubulin”, lower panel). **B.** Western blot analysis of two independent paired samples of normal (N) and tumour (T) tissues (“d” and “f”) together with the lysate from breast cancer cell line MCF7 (far right) performed as described above. **C.** Western blot analysis of lysate from breast cancer cell line MCF7 and immortalized breast cells 226LDM probed with the pre-screened anti-CTCF polyclonal antibody recognising CTCF180 and CTCF130. **D.** Western blot analysis of lysates from control and hydroxyurea/nocodazole treated 226LDM cells 226LDM cells were cell-cycle arrested by addition of 100mM hydroxyurea and 500ng/ml nocodazole in their culture medium (as described in the Supplementary materials and methods section) probed with the pre-screened anti-CTCF polyclonal antibody recognising CTCF180 and CTCF130. The development of the membranes was performed with the UptiLight™ chemiluminescence substrate. Tubulin was used as a loading control. Positions of CTCF-180, CTCF-130and tubulin are indicated. “N”- and “T” refer to normal and tumour breast tissues. **E.**Quantification of the gel from panel D performed using ImageJ [2] following the standard instructions provided by the developers. In particular, the intensities of all bands were normalised by that of the corresponding tubulin band, and then normalised by the intensity of the band of CTCF130 in control. Following normalisation the value of CTCF130 in control was designated as 1.

**Supplementary Figure S2.**
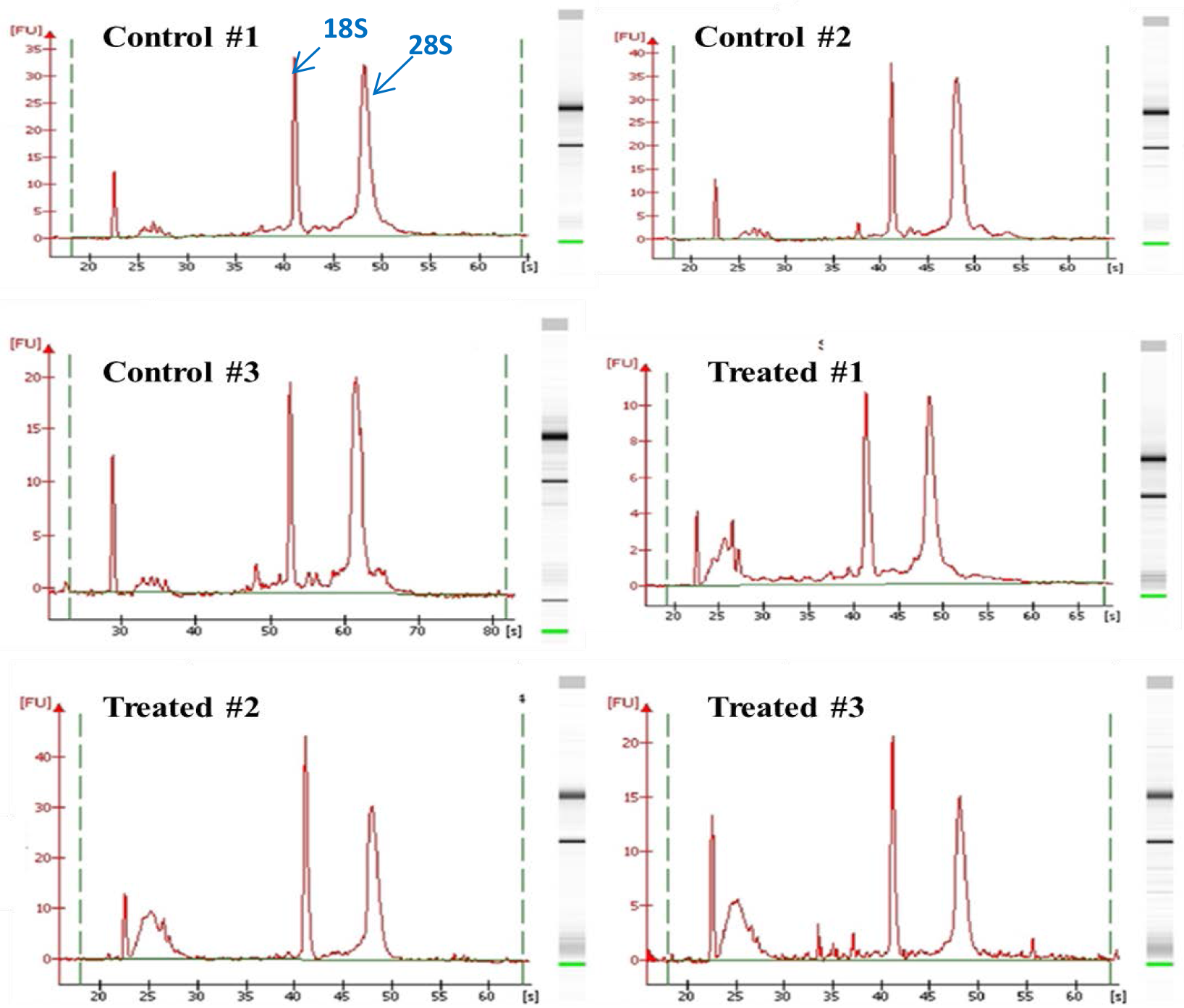
Quality control of RNA isolated from control and treated 226LDM cells. Total RNA was isolated from control and treated 226LDM cells, each group in triplicates. To confirm the integrity of the content, the samples were run on a microfluidic chip using the Agilent Bioanalyzer 2100. The 28S and 18S ribosomal subunits are represented by peaks on the electropherograph and by bands on the gel placed on the right side of each graph.

**Supplementary Figure S3.**
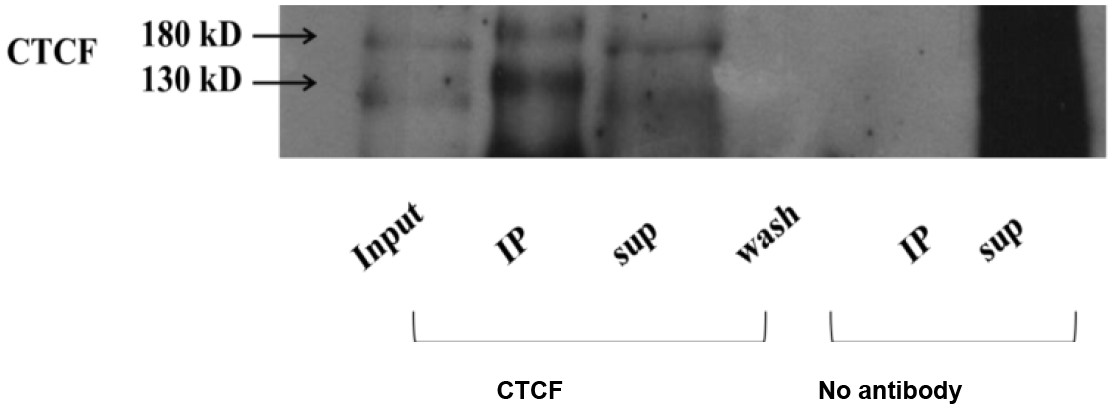
Immunoprecipitation of CTCF in 226LDM cells with the anti-CTCF polyclonal antibody: Western blot analysis. Protein extracts from untreated 226LDM cells were used in a series of protein immunoprecipitation experiments to confirm that both forms of CTCF (CTCF130 and CTCF180) can be immunoprecipitated with the selected anti-CTCF antibody (experimental details are described in the “Supplemental Materials and Methods” section). The proteins were resolved by SDS-PAGE, blotted, and probed with the selected anti-CTCF antibody. The visualization of the signal was performed using UptiLight™. Arrows indicate the positions of the two CTCF forms. Keys: Input: Pre-cleared and pre-blocked extracts (20 μl) from 226LDM cells used for the immunoprecipitation experiments. IP: Immunoprecipitated proteins (5 μl) from lysed sepharose beads. Sup: Supernatant material (20 μl) collected after centrifugation of beads with the immunoprecipitated proteins. Wash: Material (20 μl) from the wash with the immunoprecipitation buffer (BF1+BF2). No antibody: samples from the experiments performed using the same methods but without the selected CTCF antibody (used as control for the experiment.) No antibody IP: Proteins (5 μl) from lysed sepharose beads. Sup: Supernatant material (20 μl) collected after centrifugation of beads with the pre-cleared and pre-blocked extracts protein extracts.

**Supplementary Figure S4.**
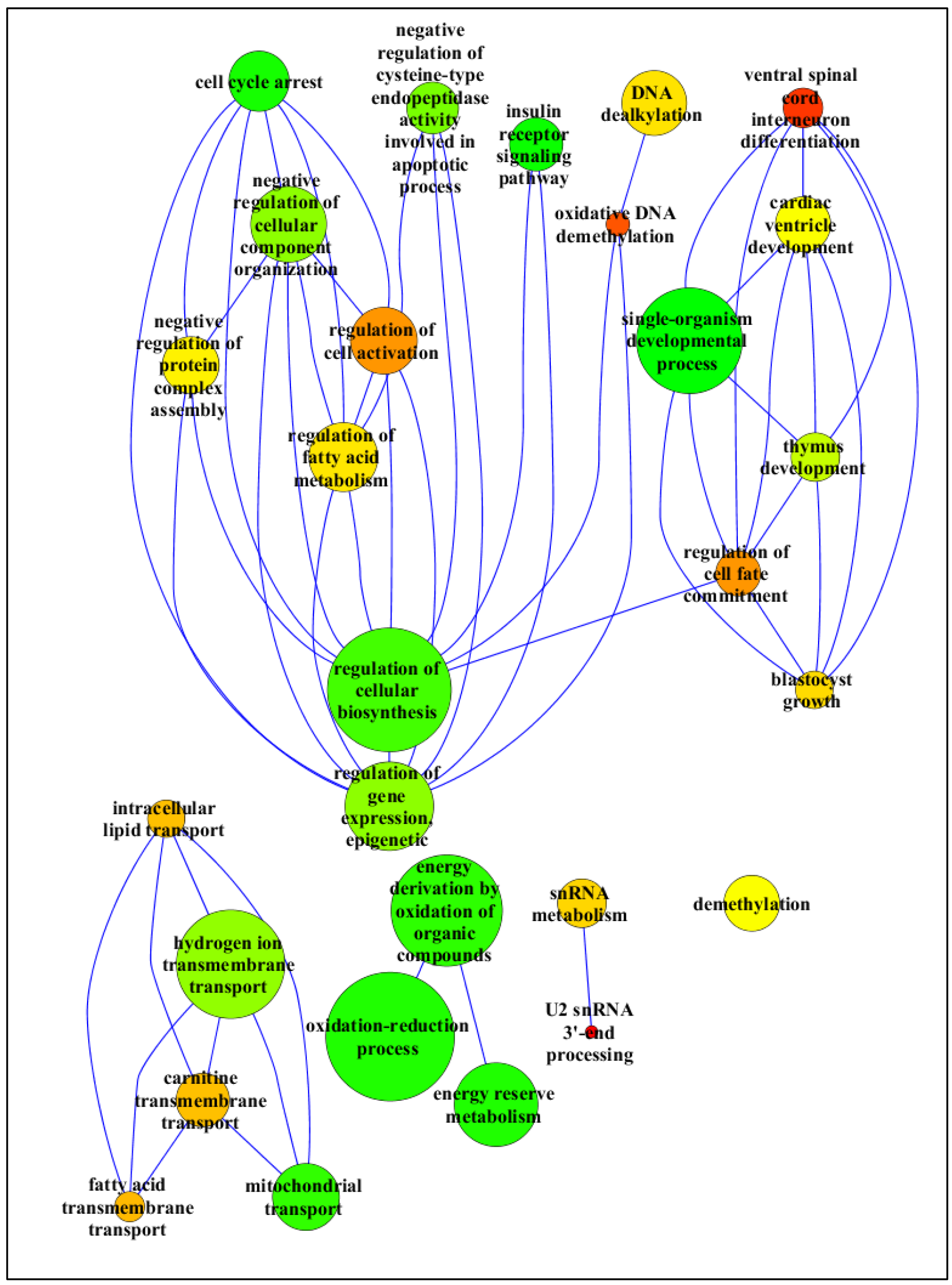
Gene ontology analysis for the genes with up-regulated expression in cell-cycle blocked 226LDM cells treated with hydroxyurea and nocodazole. Differential expression analysis was performed on control and cell-cycle blocked 226LDM cells with the DESEQ package. A gene ontology analysis was performed with the ensuing ranked list of significantly up-regulated genes. GO annotation terms were assigned to each gene and statistical Fanalysis and clustering of these terms was performed by the REViGO web server. The ontology relationships between the up-regulated genes are shown in the graph with nodes representing biological processes connected in a parent-child manner with edges. The size of the nodes represents the log-scales size of terms falling into the biological process described in the node label. The color of each node varies according to the p-value (from red to green, red representing the lowest p-value). The editing of the graph was performed with the CytoScape V3.2.1 software.

**Supplementary Figure S5.**
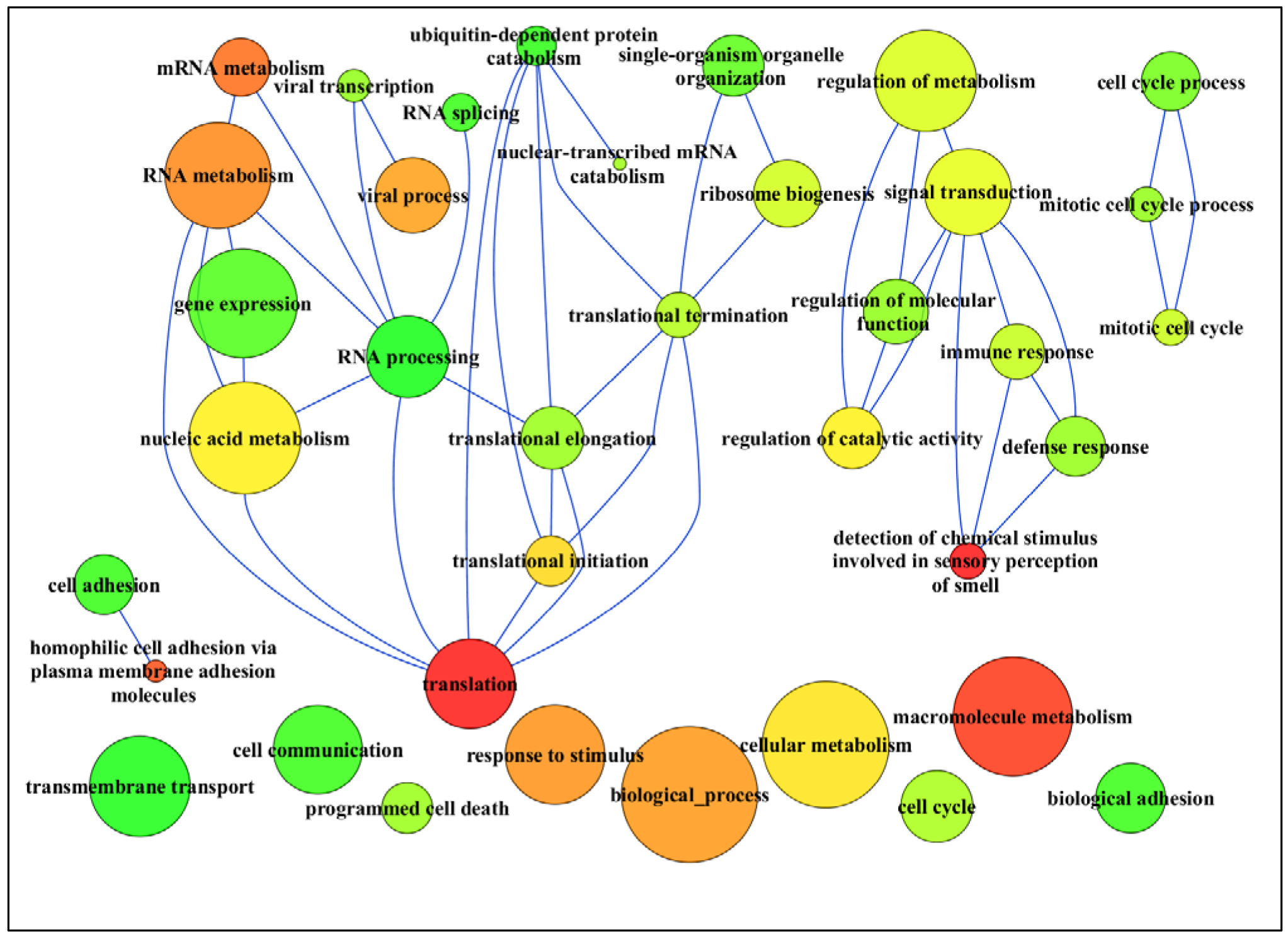
Gene ontology analysis for the genes with down-regulated expression in cell-cycle arrested 226LDM cells treated with hydroxyurea and nocodazole. Differential expression analysis was performed on control and cell-cycle blocked 226LDM cells with the DESEQ package. A gene ontology analysis was performed with the ensuing ranked list of significantly down-regulated genes. GO annotation terms were assigned to each gene and statistical analysis and clustering of these terms was performed by the REViGO web server. The ontology relationships between the down-regulated genes are shown in the graph with nodes representing biological processes connected in a parent-child manner with edges. The size of the nodes represents the log_size of terms falling into the biological process described in the node label. The color of each node varies according to the p-value (from red to green, red representing the lowest p-value). The editing of the graph was performed with the CytoScape V3.2.1 software.

**Supplementary Figure S6.**
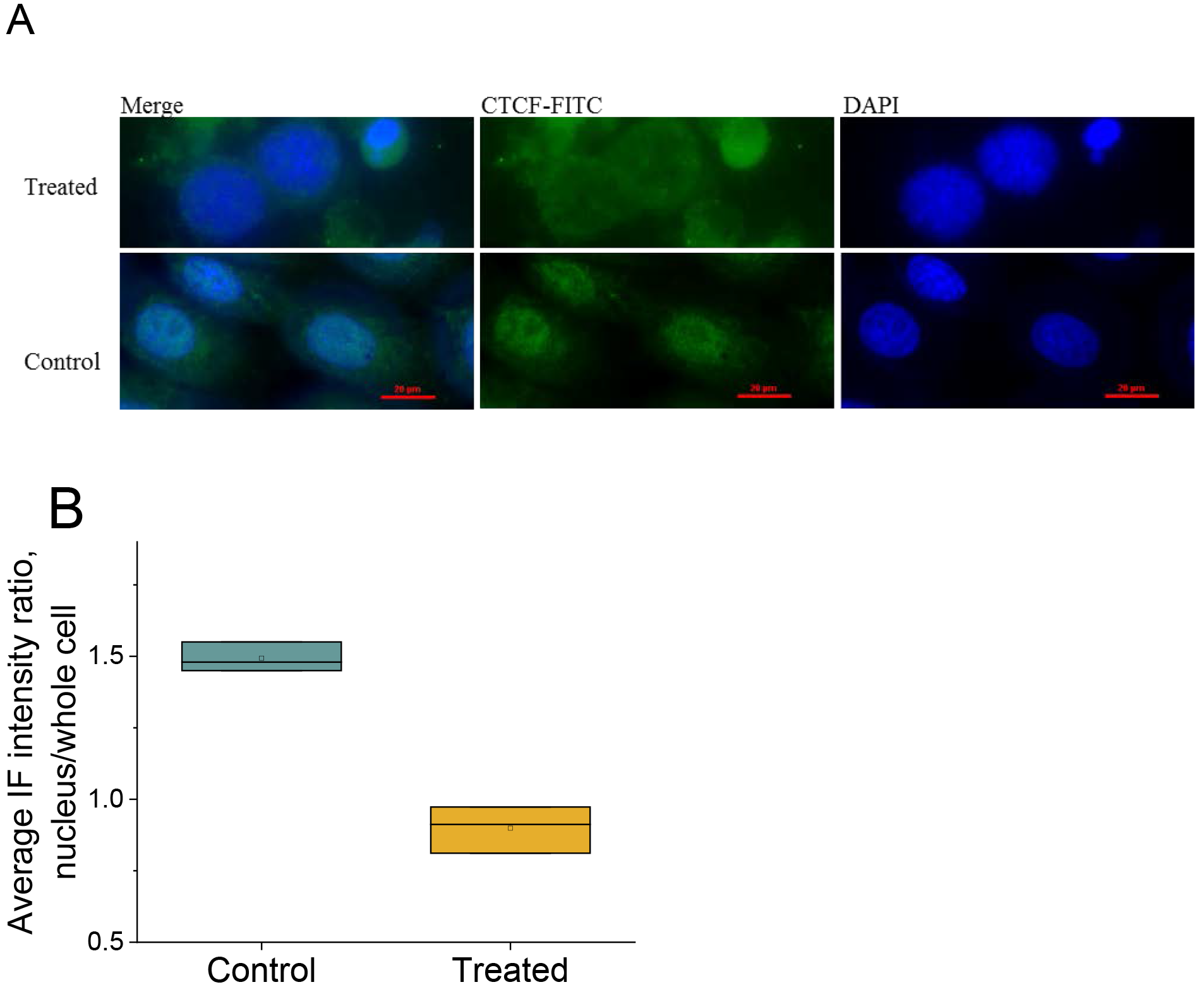
Localization profile of CTCF in control and treated 226LDM cells. **A)** CTCF immunofluorescence staining of control (untreated) and treated 226LDM cells using the polyclonal anti-CTCF antibody that recognizes both CTCF130 and CTCF180. In control cells the CTCF signal (green) appears *predominantly* diffused in the nuclear area. In treated cells staining is mostly detected in the cytoplasm. Blue colour shows DAPI staining of the nucleus. **B)** Average CTCF immunofluorescence (IF) signal intensity ratio between the nucleus and the whole cell calculated separately for the treatment and control conditions. The quantification was performed using ImageJ [2] using three individual cells (N=3) on the same slide. All ratios of intensities were normalised by that in the control. The box height shows the standard deviation calculated based on three independent measurements (three cells). The horizontal line inside the box shows the median values. Keys: “CTCF-FITC” – Staining of CTCF and with secondary antibodies conjugated with a fluorescent dyes, (e.g. FITC. “DAPI” – visualisation of nuclei with the DNA binding dye, DAPI (4’,6-diamidino-2-phenylindole, dilactate). “Merge” – overlay of CTCF-FITC and DAPI fluorescent images.

**Supplementary Figure S7.**
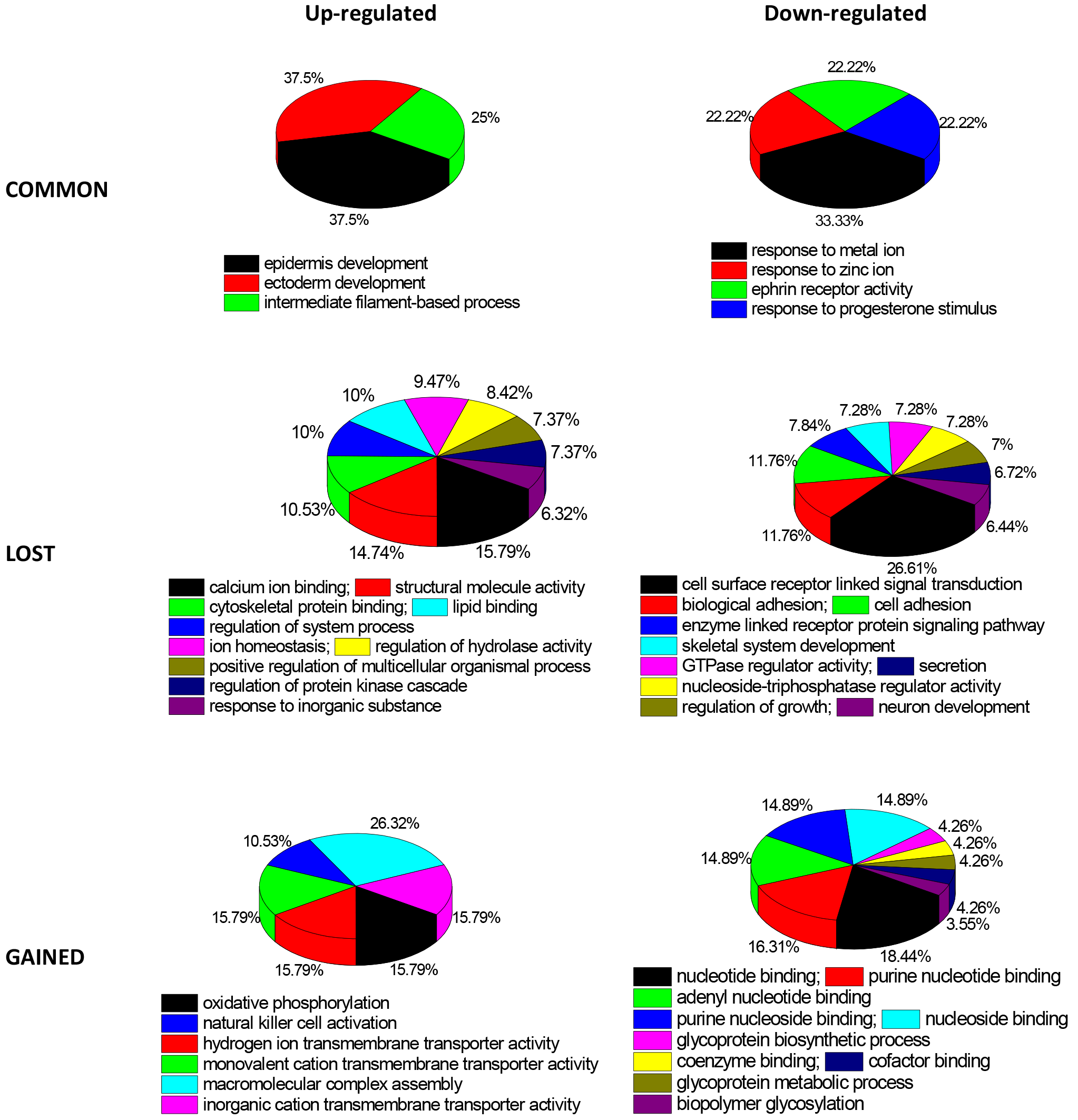
Relative enrichment of gene ontology terms of the genes whose promoters contains CTCF sites within the interval [−10,000, +1000] around CTCF. The functional classification was performed using the same datasets as in Figure 1E.

**Supplementary Figure S8.**
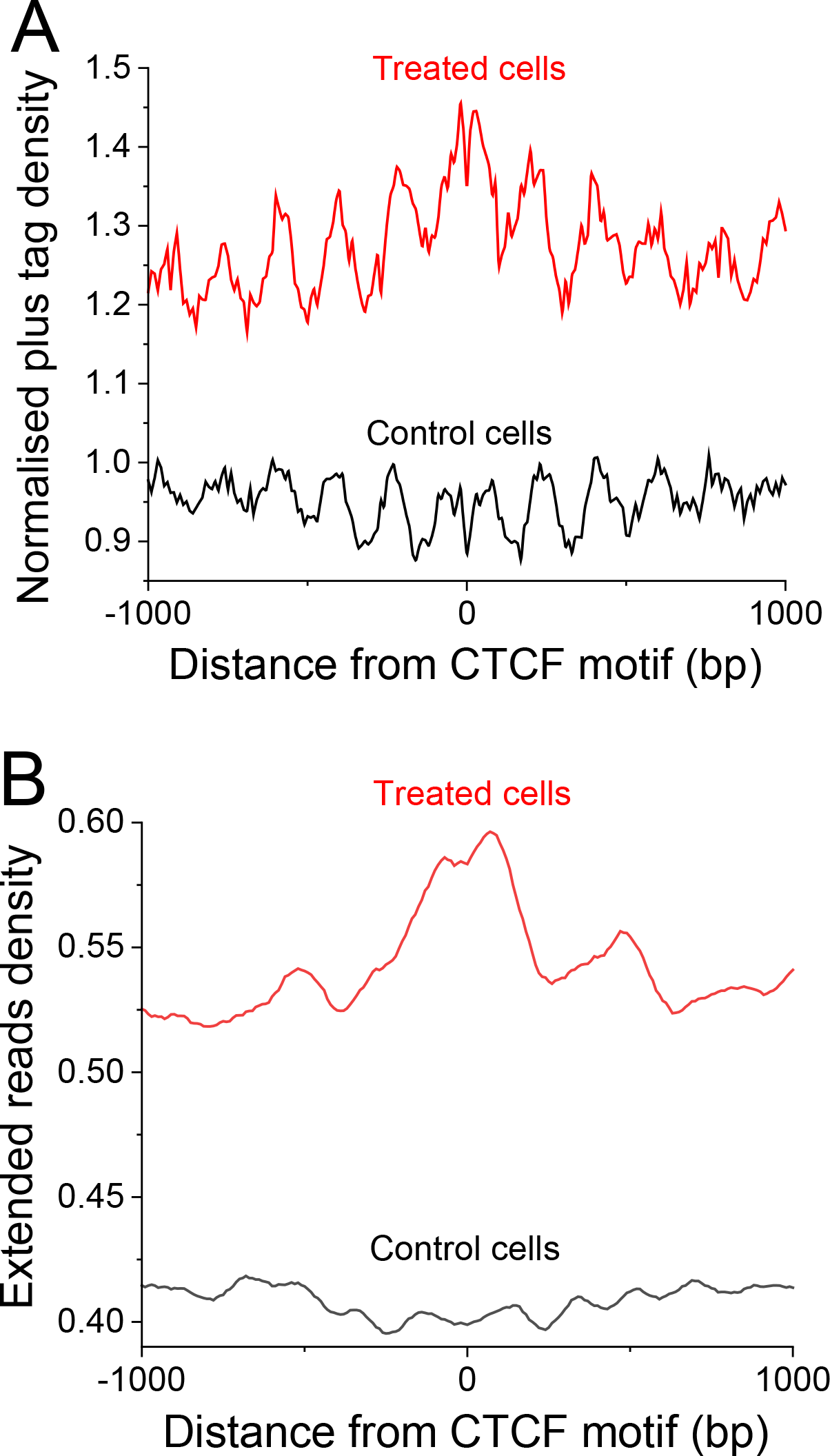
Input tag read density changes reflect nucleosome repositioning near lost CTCF sites. (A) Average profile of tag density from Input (chromatin sonication without added antibody) around the centres of CTCF motifs inside lost CTCF peaks. Each peak corresponds to the preferred individual nucleosome location. The oscillation of peaks corresponds to the nucleosome repeat length. The profile obtained after the HOMER calculation was re-normalised by dividing each line by 0.002. (B) Aggregate nucleosome occupancy profile around the centres of CTCF binding motifs inside lost CTCF peaks, calculated based on the read density from panel (A). The calculation was performed using the default HOMER procedure, which includes extending the raw tags to achieve the average DNA fragment length estimated for a given sample. The original HOMER normalisation with respect to genome average is kept. In this normalisation values above 1 mean higher than genome-average, and below 1 mean lower than genome-average. Black lines – control (untreated) cells; red lines – treated cells.

**Supplementary Figure S9.**
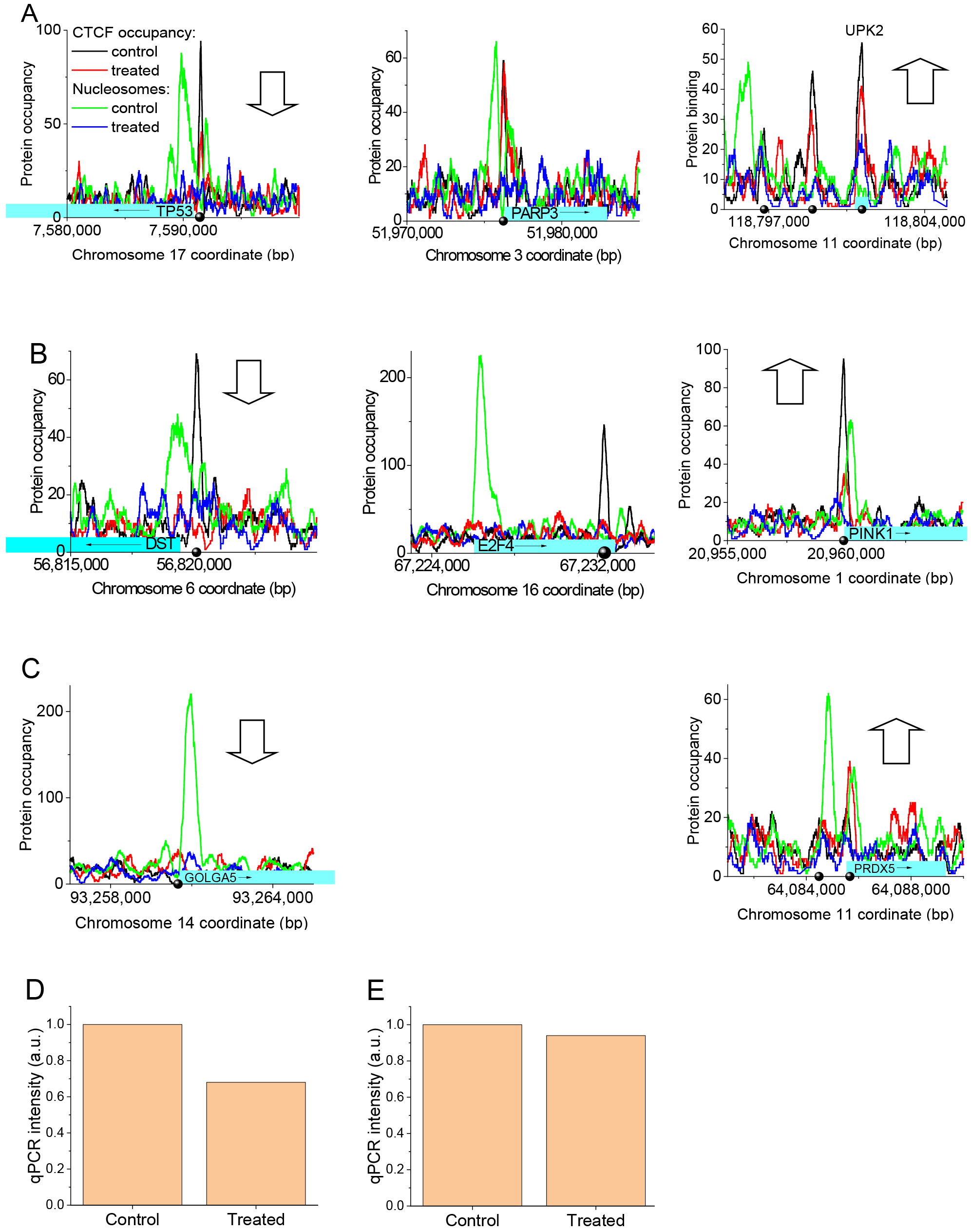
Disappearance of the chromatin density peak near CTCF upon cell treatment. (A-C) Occupancy of CTCF and the corresponding ChIP-seq Input at exemplary promoters that contain common (A), lost (B) and gained (C) CTCF. (D-E) qPCR enrichment measurement at the summit of the chromatin peak positions given by the green line in panel A using Input and no-antibody control for promoters PARP3 (D) and TP53 (E). qPCR normalisation of Input samples was performed as described in Supplementary Methods.

**Supplementary Figure S10.**
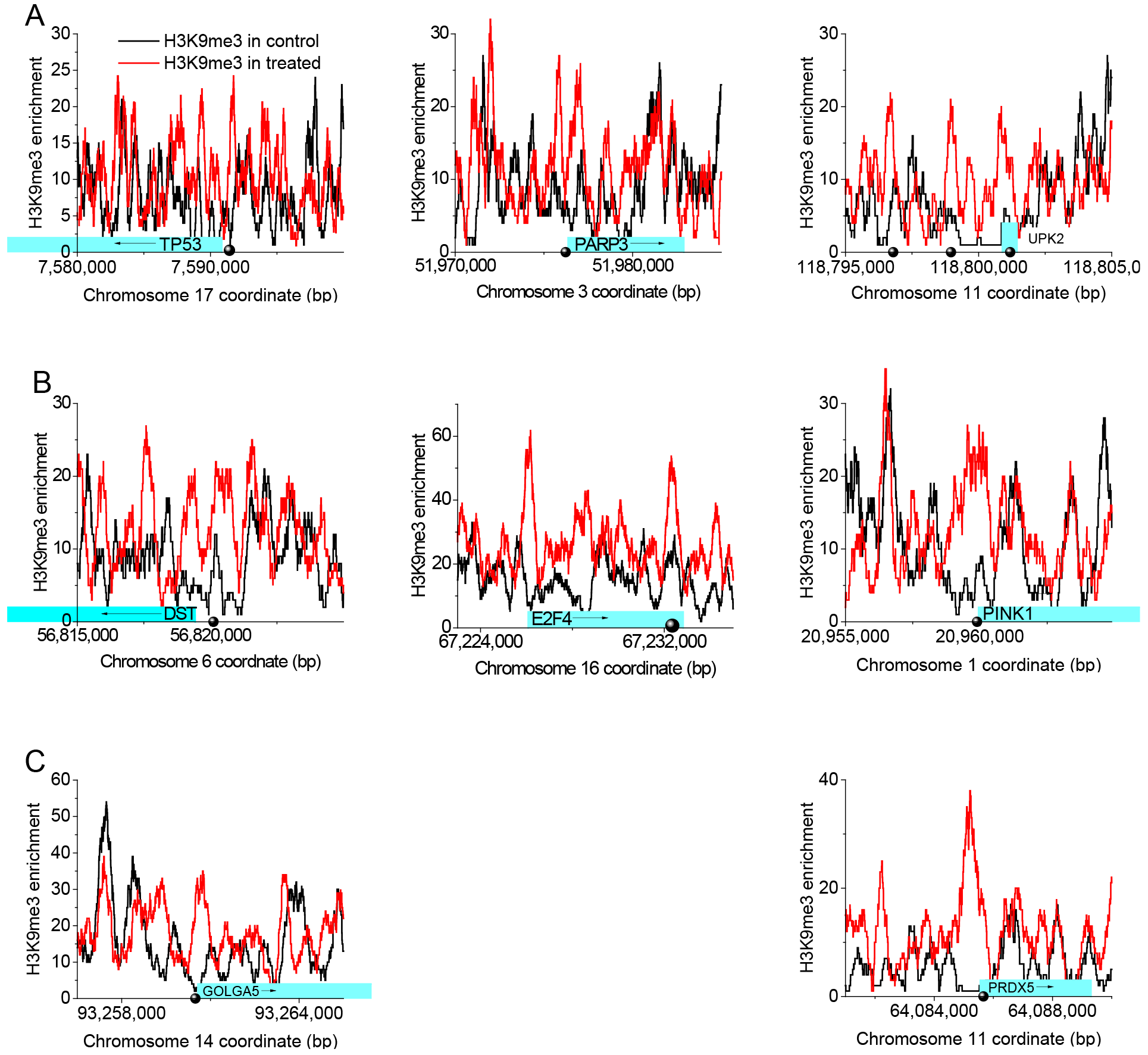
Enrichment of H3K9me3 in control (black line) and treated cells (red line) at exemplary promoters that contain common (A), lost (B) and gained (C) CTCF. The same promoters as in Figure S9 are selected.

## REFERENCES

[1] C.T. Ong, V.G. Corces, CTCF: an architectural protein bridging genome topology and function, Nat Rev Genet, 15 (2014) 234–246.

[2] G. Bonora, K. Plath, M. Denholtz, A mechanistic link between gene regulation and genome architecture in mammalian development, Current opinion in genetics & development, 27 (2014) 92–101.

[3] T. Ali, R. Renkawitz, M. Bartkuhn, Insulators and domains of gene expression, Curr Opin Genet Dev, 37 (2016) 17–26.

[4] M. Merkenschlager, D.T. Odom, CTCF and cohesin: linking gene regulatory elements with their targets, Cell, 152 (2013) 1285–1297.

[5] S.J. Holwerda, W. de Laat, CTCF: the protein, the binding partners, the binding sites and their chromatin loops, Philos Trans R Soc Lond B Biol Sci, 368 (2013) 20120369.

[6] L.L.P. Hanssen, M.T. Kassouf, A.M. Oudelaar, D. Biggs, C. Preece, D.J. Downes, M. Gosden, J.A. Sharpe, J.A. Sloane-Stanley, J.R. Hughes, B. Davies, D.R. Higgs, Tissue-specific CTCF-cohesin-mediated chromatin architecture delimits enhancer interactions and function in vivo, Nat Cell Biol, 19 (2017) 952–961.

[7] A.C. Mullen, A.S. Hutchins, A.V. Villarino, H.W. Lee, F.A. High, N. Cereb, S.Y. Yang, X. Hua, S.L. Reiner, Cell cycle controlling the silencing and functioning of mammalian activators, Curr Biol, 11 (2001) 1695–1699.

[8] R. Ohlsson, V. Lobanenkov, E. Klenova, Does CTCF mediate between nuclear organization and gene expression?, Bioessays, 32 (2010) 37–50.

[9] D. Hnisz, D.S. Day, R.A. Young, Insulated Neighborhoods: Structural and Functional Units of Mammalian Gene Control, Cell, 167 (2016) 1188–1200.

[10] A. Mishra, R.D. Hawkins, Three-dimensional genome architecture and emerging technologies: looping in disease, Genome Med, 9 (2017) 87.

[11] A. Canela, Y. Maman, S. Jung, N. Wong, E. Callen, A. Day, K.R. Kieffer-Kwon, A. Pekowska, H. Zhang, S.S.P. Rao, S.C. Huang, P.J. McKinnon, P.D. Aplan, Y. Pommier, E.L. Aiden, R. Casellas, A. Nussenzweig, Genome Organization Drives Chromosome Fragility, Cell, 170 (2017) 507–521.

[12] Y. Lu, G. Shan, J. Xue, C. Chen, C. Zhang, Defining the multivalent functions of CTCF from chromatin state and three-dimensional chromatin interactions, Nucleic acids research, 44 (2016) 6200–6212.

[13] H. Wang, M.T. Maurano, H. Qu, K.E. Varley, J. Gertz, F. Pauli, K. Lee, T. Canfield, M. Weaver, R. Sandstrom, R.E. Thurman, R. Kaul, R.M. Myers, J.A. Stamatoyannopoulos, Widespread plasticity in CTCF occupancy linked to DNA methylation, Genome Res, 22 (2012) 1680–1688.

[14] W.L. Kraus, M.O. Hottiger, PARP-1 and gene regulation: progress and puzzles, Molecular aspects of medicine, 34 (2013) 1109–1123.

[15] C. Thomas, A.V. Tulin, Poly-ADP-ribose polymerase: machinery for nuclear processes, Molecular aspects of medicine, 34 (2013) 1124–1137.

[16] B. Golia, H.R. Singh, G. Timinszky, Poly-ADP-ribosylation signaling during DNA damage repair, Front Biosci (Landmark Ed), 20 (2015) 440–457.

[17] I. Robert, O. Karicheva, B. Reina San Martin, V. Schreiber, F. Dantzer, Functional aspects of PARylation in induced and programmed DNA repair processes: preserving genome integrity and modulating physiological events, Molecular aspects of medicine, 34 (2013) 1138–1152.

[18] A. Burkle, Poly(ADP-ribosyl)ation: a posttranslational protein modification linked with genome protection and mammalian longevity, Biogerontology, 1 (2000) 41–46.

[19] M. Li, X. Yu, The role of poly(ADP-ribosyl)ation in DNA damage response and cancer chemotherapy, Oncogene, 34 (2015) 3349–3356.

[20] P. Caiafa, J. Zlatanova, CCCTC-binding factor meets poly(ADP-ribose) polymerase-1, Journal of cellular physiology, 219 (2009) 265–270.

[21] E. Klenova, R. Ohlsson, Poly(ADP-ribosyl)ation and epigenetics. Is CTCF PARt of the plot?, Cell cycle, 4 (2005) 96–101.

[22] D. Han, Q. Chen, J. Shi, F. Zhang, X. Yu, CTCF participates in DNA damage response via poly(ADP-ribosyl)ation, Sci Rep, 7 (2017) 43530.

[23] D. Farrar, S. Rai, I. Chernukhin, M. Jagodic, Y. Ito, S. Yammine, R. Ohlsson, A. Murrell, E. Klenova, Mutational Analysis of the Poly(ADP-Ribosyl)ation Sites of the Transcription Factor CTCF Provides an Insight into the Mechanism of Its Regulation by Poly(ADP-Ribosyl)ation, Molecular and cellular biology, 30 (2010) 1199–1216.

[24] T. Guastafierro, B. Cecchinelli, M. Zampieri, A. Reale, G. Riggio, O. Sthandier, G. Zupi, L. Calabrese, P. Caiafa, CCCTC-binding factor activates PARP-1 affecting DNA methylation machinery, J Biol Chem, 283 (2008) 21873–21880.

[25] N. Nalabothula, T. Al-Jumaily, A.M. Eteleeb, R.M. Flight, S. Xiaorong, H. Moseley, E.C. Rouchka, Y.N. Fondufe-Mittendorf, Genome-Wide Profiling of PARP1 Reveals an Interplay with Gene Regulatory Regions and DNA Methylation, PloS one, 10 (2015) e0135410.

[26] H. Zhao, E.G. Sifakis, N. Sumida, L. Millan-Arino, B.A. Scholz, J.P. Svensson, X. Chen, A.L. Ronnegren, C.D. Mallet de Lima, F.S. Varnoosfaderani, C. Shi, O. Loseva, S. Yammine, M. Israelsson, L.S. Rathje, B. Nemeti, E. Fredlund, T. Helleday, M.P. Imreh, A. Gondor, PARP1- and CTCF-Mediated Interactions between Active and Repressed Chromatin at the Lamina Promote Oscillating Transcription, Mol Cell, 59 (2015) 984–997.

[27] W. Yu, V. Ginjala, V. Pant, I. Chernukhin, J. Whitehead, F. Docquier, D. Farrar, G. Tavoosidana, R. Mukhopadhyay, C. Kanduri, M. Oshimura, A.P. Feinberg, V. Lobanenkov, E. Klenova, R. Ohlsson, Poly(ADP-ribosyl)ation regulates CTCF-dependent chromatin insulation, Nat Genet, 36 (2004) 1105–1110.

[28] V. Torrano, J. Navascues, F. Docquier, R. Zhang, L.J. Burke, I. Chernukhin, D. Farrar, J. Leon, M.T. Berciano, R. Renkawitz, E. Klenova, M. Lafarga, M.D. Delgado, Targeting of CTCF to the nucleolus inhibits nucleolar transcription through a poly(ADP-ribosyl)ation-dependent mechanism, J Cell Sci, 119 (2006) 1746–1759.

[29] F. Docquier, G.X. Kita, D. Farrar, P. Jat, M. O’Hare, I. Chernukhin, S. Gretton, A. Mandal, L. Alldridge, E. Klenova, Decreased poly(ADP-ribosyl)ation of CTCF, a transcription factor, is associated with breast cancer phenotype and cell proliferation, Clinical Cancer Research, 15 (2009) 5762–5771.

[30] I. Pavlaki, Role of CTCF Poly(ADP-ribosyl)ation in the regulation of cellular functions, School of Biological Sciences, University of Essex, UK, Colchester, Essex, UK, 2016.

[31] A. Chrambach, D. Rodbard, Polyacrylamide gel electrophoresis, Science, 172 (1971) 440–451.

[32] H. Towbin, T. Staehelin, J. Gordon, Electrophoretic transfer of proteins from polyacrylamide gels to nitrocellulose sheets: procedure and some applications, Proc Natl Acad Sci U S A, 76 (1979) 4350–4354.

[33] B. Kaboord, M. Perr, Isolation of proteins and protein complexes by immunoprecipitation, Methods in molecular biology, 424 (2008) 349–364.

[34] B. Langmead, C. Trapnell, M. Pop, S.L. Salzberg, Ultrafast and memory-efficient alignment of short DNA sequences to the human genome, Genome Biol, 10 (2009) R25.

[35] Y. Zhang, T. Liu, C.A. Meyer, J. Eeckhoute, D.S. Johnson, B.E. Bernstein, C. Nusbaum, R.M. Myers, M. Brown, W. Li, X.S. Liu, Model-based Analysis of ChIP-Seq (MACS), Genome Biol, 9 (2008) R137.

[36] A.R. Quinlan, BEDTools: The Swiss-Army Tool for Genome Feature Analysis, Curr Protoc Bioinformatics, 47 (2014) 11 12 11–34.

[37] Y. Vainshtein, K. Rippe, V.B. Teif, NucTools: analysis of chromatin feature occupancy profiles from high-throughput sequencing data, BMC Genomics, 18 (2017) 158.

[38] V.B. Teif, D.A. Beshnova, Y. Vainshtein, C. Marth, J.P. Mallm, T. Höfer, K. Rippe, Nucleosome repositioning links DNA (de)methylation and differential CTCF binding during stem cell development, Genome Res, 24 (2014) 1285–1295.

[39] S. Heinz, C. Benner, N. Spann, E. Bertolino, Y.C. Lin, P. Laslo, J.X. Cheng, C. Murre, H. Singh, C.K. Glass, Simple combinations of lineage-determining transcription factors prime cis-regulatory elements required for macrophage and B cell identities, Mol Cell, 38 (2010) 576–589.

[40] A. Mathelier, O. Fornes, D.J. Arenillas, C.Y. Chen, G. Denay, J. Lee, W. Shi, C. Shyr, G. Tan, R. Worsley-Hunt, A.W. Zhang, F. Parcy, B. Lenhard, A. Sandelin, W.W. Wasserman, JASPAR 2016: a major expansion and update of the open-access database of transcription factor binding profiles, Nucleic Acids Res, 44 (2016) D110–115.

[41] J.A. Castro-Mondragon, S. Jaeger, D. Thieffry, M. Thomas-Chollier, J. van Helden, RSAT matrix-clustering: dynamic exploration and redundancy reduction of transcription factor binding motif collections, Nucleic Acids Res, 45 (2017) e119.

[42] G. Dennis, Jr., B.T. Sherman, D.A. Hosack, J. Yang, W. Gao, H.C. Lane, R.A. Lempicki, DAVID: Database for Annotation, Visualization, and Integrated Discovery, Genome Biol, 4 (2003) P3.

[43] F. Supek, M. Bosnjak, N. Skunca, T. Smuc, REVIGO summarizes and visualizes long lists of gene ontology terms, PLoS One, 6 (2011) e21800.

[44] P. Shannon, A. Markiel, O. Ozier, N.S. Baliga, J.T. Wang, D. Ramage, N. Amin, B. Schwikowski, T. Ideker, Cytoscape: a software environment for integrated models of biomolecular interaction networks, Genome Res, 13 (2003) 2498–2504.

[45] H. Mi, X. Huang, A. Muruganujan, H. Tang, C. Mills, D. Kang, P.D. Thomas, PANTHER version 11: expanded annotation data from Gene Ontology and Reactome pathways, and data analysis tool enhancements, Nucleic Acids Res, 45 (2017) D183–D189.

[46] E. Eisenberg, E.Y. Levanon, Human housekeeping genes, revisited, Trends Genet, 29 (2013) 569–574.

[47] M.G. van Oijen, R.H. Medema, P.J. Slootweg, G. Rijksen, Positivity of the proliferation marker Ki-67 in noncycling cells, American journal of clinical pathology, 110 (1998) 24–31.

[48] V.B. Teĭf, A.V. Shkrobkov, V.P. Egorova, V.I. Krot, [Nucleosomes in gene regulation: theoretical approaches], Mol Biol (Mosk), 46 (2012) 3–13.

[49] E.P. Nora, A. Goloborodko, A.L. Valton, J.H. Gibcus, A. Uebersohn, N. Abdennur, J. Dekker, L.A. Mirny, B.G. Bruneau, Targeted Degradation of CTCF Decouples Local Insulation of Chromosome Domains from Genomic Compartmentalization, Cell, 169 (2017) 930–944.

[50] B.D. Pope, T. Ryba, V. Dileep, F. Yue, W. Wu, O. Denas, D.L. Vera, Y. Wang, R.S. Hansen, T.K. Canfield, R.E. Thurman, Y. Cheng, G. Gulsoy, J.H. Dennis, M.P. Snyder, J.A. Stamatoyannopoulos, J. Taylor, R.C. Hardison, T. Kahveci, B. Ren, D.M. Gilbert, Topologically associating domains are stable units of replication-timing regulation, Nature, 515 (2014) 402–405.

[51] V.B. Teif, F. Erdel, D.A. Beshnova, Y. Vainshtein, J.P. Mallm, K. Rippe, Taking into account nucleosomes for predicting gene expression, Methods, 62 (2013) 26–38.

[52] T.H. Kim, Z.K. Abdullaev, A.D. Smith, K.A. Ching, D.I. Loukinov, R.D. Green, M.Q. Zhang, V.V. Lobanenkov, B. Ren, Analysis of the vertebrate insulator protein CTCF-binding sites in the human genome, Cell, 128 (2007) 1231–1245.

[53] J.D. Ziebarth, A. Bhattacharya, Y. Cui, CTCFBSDB 2.0: a database for CTCF-binding sites and genome organization, Nucleic Acids Res, 41 (2013) D188–194.

[54] R.N. Plasschaert, S. Vigneau, I. Tempera, R. Gupta, J. Maksimoska, L. Everett, R. Davuluri, R. Mamorstein, P.M. Lieberman, D. Schultz, S. Hannenhalli, M.S. Bartolomei, CTCF binding site sequence differences are associated with unique regulatory and functional trends during embryonic stem cell differentiation, Nucleic acids research, 42 (2014) 774–789.

[55] K. Essien, S. Vigneau, S. Apreleva, L.N. Singh, M.S. Bartolomei, S. Hannenhalli, CTCF binding site classes exhibit distinct evolutionary, genomic, epigenomic and transcriptomic features, Genome Biol, 10 (2009) R131.

[56] S. Cuddapah, R. Jothi, D.E. Schones, T.Y. Roh, K. Cui, K. Zhao, Global analysis of the insulator binding protein CTCF in chromatin barrier regions reveals demarcation of active and repressive domains, Genome Res, 19 (2009) 24–32.

[57] Y. Fu, M. Sinha, C.L. Peterson, Z. Weng, The insulator binding protein CTCF positions 20 nucleosomes around its binding sites across the human genome, PLoS Genetics, 4 (2008) e1000138.

[58] V.B. Teif, Y. Vainshtein, M. Caudron-Herger, J.P. Mallm, C. Marth, T. Höfer, K. Rippe, Genome-wide nucleosome positioning during embryonic stem cell development, Nat Struct Mol Biol, 19 (2012) 1185–1192.

[59] D.A. Beshnova, A.G. Cherstvy, Y. Vainshtein, V.B. Teif, Regulation of the nucleosome repeat length in vivo by the DNA sequence, protein concentrations and long-range interactions, PLoS Comput Biol, 10 (2014) e1003698.

[60] I. Kremsky, N. Bellora, E. Eyras, A Quantitative Profiling Tool for Diverse Genomic Data Types Reveals Potential Associations between Chromatin and Pre-mRNA Processing, PLoS One, 10 (2015) e0132448.

[61] Y.B. Schwartz, T.G. Kahn, V. Pirrotta, Characteristic low density and shear sensitivity of crosslinked chromatin containing polycomb complexes, Molecular and cellular biology, 25 (2005) 432–439.

[62] J.S. Becker, R.L. McCarthy, S. Sidoli, G. Donahue, K.E. Kaeding, Z. He, S. Lin, B.A. Garcia, K.S. Zaret, Genomic and Proteomic Resolution of Heterochromatin and Its Restriction of Alternate Fate Genes, Mol Cell, 68 (2017) 1023–1037.

[63] M.S. Poptsova, I.A. Il’icheva, D.Y. Nechipurenko, L.A. Panchenko, M.V. Khodikov, N.Y. Oparina, R.V. Polozov, Y.D. Nechipurenko, S.L. Grokhovsky, Non-random DNA fragmentation in next-generation sequencing, Sci Rep, 4 (2014) 4532.

[64] A. Lazarovici, T. Zhou, A. Shafer, A.C. Dantas Machado, T.R. Riley, R. Sandstrom, P.J. Sabo, Y. Lu, R. Rohs, J.A. Stamatoyannopoulos, H.J. Bussemaker, Probing DNA shape and methylation state on a genomic scale with DNase I, Proc Natl Acad Sci U S A, 110 (2013) 6376–6381.

[65] S.L. Grokhovsky, I.A. Il’icheva, D.Y. Nechipurenko, M.V. Golovkin, L.A. Panchenko, R.V. Polozov, Y.D. Nechipurenko, Sequence-specific ultrasonic cleavage of DNA, Biophys J, 100 (2011) 117–125.

[66] D. Tillo, N. Kaplan, I.K. Moore, Y. Fondufe-Mittendorf, A.J. Gossett, Y. Field, J.D. Lieb, J. Widom, E. Segal, T.R. Hughes, High nucleosome occupancy is encoded at human regulatory sequences, PLoS ONE, 5 (2010) e9129.

[67] M.T. Hayashi, J. Karlseder, DNA damage associated with mitosis and cytokinesis failure, Oncogene, 32 (2013) 4593–4601.

[68] I. Chaudhury, D.M. Koepp, Recovery from the DNA Replication Checkpoint, Genes (Basel), 7 (2016) 94.

[69] H.E. Bryant, E. Petermann, N. Schultz, A.S. Jemth, O. Loseva, N. Issaeva, F. Johansson, S. Fernandez, P. McGlynn, T. Helleday, PARP is activated at stalled forks to mediate Mre11-dependent replication restart and recombination, The EMBO journal, 28 (2009) 2601–2615.

[70] J. Yelamos, J. Farres, L. Llacuna, C. Ampurdanes, J. Martin-Caballero, PARP-1 and PARP-2: New players in tumour development, Am J Cancer Res, 1 (2011) 328–346.

[71] K.L. MacQuarrie, A.P. Fong, R.H. Morse, S.J. Tapscott, Genome-wide transcription factor binding: beyond direct target regulation, Trends Genet, 27 (2011) 141–148.

[72] M. Tark-Dame, H. Jerabek, E.M. Manders, I.M. van der Wateren, D.W. Heermann, R. van Driel, Depletion of the chromatin looping proteins CTCF and cohesin causes chromatin compaction: insight into chromatin folding by polymer modelling, PLoS Comput Biol, 10 (2014) e1003877.

[73] C. Lopes Novo, P.J. Rugg-Gunn, Chromatin organization in pluripotent cells: emerging approaches to study and disrupt function, Briefings in functional genomics, 15 (2016) 305–314.

[74] N.S. Benabdallah, I. Williamson, R.S. Illingworth, S. Boyle, G.R. Grimes, P. Therizols, W. Bickmore, PARP mediated chromatin unfolding is coupled to long-range enhancer activation, bioRxiv, (2017) 155325.

[75] R.K. Auerbach, G. Euskirchen, J. Rozowsky, N. Lamarre-Vincent, Z. Moqtaderi, P. Lefrancois, K. Struhl, M. Gerstein, M. Snyder, Mapping accessible chromatin regions using Sono-Seq, Proc Natl Acad Sci U S A, 106 (2009) 14926–14931.

[76] L. Teytelman, D.M. Thurtle, J. Rine, A. van Oudenaarden, Highly expressed loci are vulnerable to misleading ChIP localization of multiple unrelated proteins, Proc Natl Acad Sci U S A, 110 (2013) 18602–18607.

[77] L. Teytelman, B. Ozaydin, O. Zill, P. Lefrancois, M. Snyder, J. Rine, M.B. Eisen, Impact of chromatin structures on DNA processing for genomic analyses, PLoS One, 4 (2009) e6700.

[78] Y. Guo, Q. Xu, D. Canzio, J. Shou, J. Li, D.U. Gorkin, I. Jung, H. Wu, Y. Zhai, Y. Tang, Y. Lu, Y. Wu, Z. Jia, W. Li, M.Q. Zhang, B. Ren, A.R. Krainer, T. Maniatis, Q. Wu, CRISPR Inversion of CTCF Sites Alters Genome Topology and Enhancer/Promoter Function, Cell, 162 (2015) 900–910.

[79] E. de Wit, E.S. Vos, S.J. Holwerda, C. Valdes-Quezada, M.J. Verstegen, H. Teunissen, E. Splinter, P.J. Wijchers, P.H. Krijger, W. de Laat, CTCF Binding Polarity Determines Chromatin Looping, Mol Cell, 60 (2015) 676–684.

[80] A.S. Hansen, I. Pustova, C. Cattoglio, R. Tjian, X. Darzacq, CTCF and cohesin regulate chromatin loop stability with distinct dynamics, Elife, 6 (2017) e25776.

[81] D. Tempka, P. Tokarz, K. Chmielewska, M. Kluska, J. Pietrzak, Z. Rygielska, L. Virag, A. Robaszkiewicz, Downregulation of PARP1 transcription by CDK4/6 inhibitors sensitizes human lung cancer cells to anticancer drug-induced death by impairing OGG1-dependent base excision repair, Redox Biol, 15 (2018) 316–326.

[82] T. Sekiya, K. Murano, K. Kato, A. Kawaguchi, K. Nagata, Mitotic phosphorylation of CCCTC-binding factor (CTCF) reduces its DNA binding activity, FEBS Open Bio, 7 (2017) 397–404.

[83] H. Agarwal, M. Reisser, C. Wortmann, J.C.M. Gebhardt, Direct Observation of Cell-Cycle-Dependent Interactions between CTCF and Chromatin, Biophysical journal, 112 (2017) 2051–2055.

[84] L.J. Burke, R. Zhang, M. Bartkuhn, V.K. Tiwari, G. Tavoosidana, S. Kurukuti, C. Weth, J. Leers, N. Galjart, R. Ohlsson, R. Renkawitz, CTCF binding and higher order chromatin structure of the H19 locus are maintained in mitotic chromatin, EMBO J, 24 (2005) 3291–3300.

[85] W. Shen, D. Wang, B. Ye, M. Shi, Y. Zhang, Z. Zhao, A possible role of Drosophila CTCF in mitotic bookmarking and maintaining chromatin domains during the cell cycle, Biol Res, 48 (2015) 27

[86] N. Festuccia, I. Gonzalez, N. Owens, P. Navarro, Mitotic bookmarking in development and stem cells, Development, 144 (2017) 3633–3645.

[87] M. Nekrasov, J. Amrichova, B.J. Parker, T.A. Soboleva, C. Jack, R. Williams, G.A. Huttley, D.J. Tremethick, Histone H2A.Z inheritance during the cell cycle and its impact on promoter organization and dynamics, Nature structural & molecular biology, 19 (2012) 1076–1083.

## Supplemental References

[1] F. Docquier, G.X. Kita, D. Farrar, P. Jat, M. O’Hare, I. Chernukhin, S. Gretton, A. Mandal, L. Alldridge, E. Klenova, Decreased poly(ADP-ribosyl)ation of CTCF, a transcription factor, is associated with breast cancer phenotype and cell proliferation, Clinical cancer research: an official journal of the American Association for Cancer Research, 15 (2009) 5762–5771.

[2] C.T. Rueden, J. Schindelin, M.C. Hiner, B.E. DeZonia, A.E. Walter, E.T. Arena, K.W. Eliceiri, ImageJ2: ImageJ for the next generation of scientific image data, BMC Bioinformatics, 18 (2017) 529.

[3] J.A. Ramos-Vara, Technical aspects of immunohistochemistry, Vet Pathol, 42 (2005) 405–426.

[4] K.J. Livak, T.D. Schmittgen, Analysis of Relative Gene Expression Data Using Real-Time Quantitative PCR and the 2-ΔΔCT Method, Methods, 25 (2001) 402–408.

[5] V. D’Arcy, N. Pore, F. Docquier, Z.K. Abdullaev, I. Chernukhin, G.X. Kita, S. Rai, M. Smart, D. Farrar, S. Pack, V. Lobanenkov, E. Klenova, BORIS, a paralogue of the transcription factor, CTCF, is aberrantly expressed in breast tumours, British journal of cancer, 98 (2008) 571–579.

